# Turning red to green repeatedly: ancient LWS duplications drive convergent visual tuning in teleost fishes

**DOI:** 10.1101/2021.05.08.443214

**Authors:** Fabio Cortesi, Daniel Escobar Camacho, Gina Maria Sommer, Martin Luehrmann, Flurin Hidber, Xavier Deupi, Karen Carleton, Zuzana Musilova

## Abstract

Photopigments, formed by an opsin protein bound to a light-sensitive chromophore, underlie vertebrate vision. Long-wavelength–sensitive (LWS) opsins mediate red-light detection, and most teleosts retain a single functional LWS1. A shorter-shifted green-sensitive paralog (LWS2) is found only in a few lineages. By mining teleost genomes and sequencing retinal transcriptomes, we identify elopomorphs as an additional lineage retaining LWS2 (alongside characins and osteoglossomorphs), and we reveal a previously overlooked shorter-shifted paralog, LWS3, restricted to gobies (Percomorpha) and arising from an ancient duplication. Structural modeling of twelve LWS opsins reveals convergent evolution at four key amino acid sites (214, 259, 261, 269) in the retinal-binding pocket, indicating convergent substitutions in human MWS, teleost LWS2, and goby LWS3 relative to red-sensitive counterparts, consistent with repeated spectral shifts toward green wavelengths. In several lineages—including characins, mormyrids, gobies, and primates—these shorter-shifted LWS opsins have functionally replaced the canonical green-sensitive RH2 opsin. Retinal transcriptomes and in situ hybridization demonstrate variable *lws3* expression across gobies, with localization to a distinct double-cone member in *Amblygobius phalaena*, analogous to *rh2* expression in other fish species. Together, these results show that repeated convergent evolution toward green sensitivity over 500 million years involves coordinated changes at the molecular, regulatory, and functional levels, providing a striking example of multilevel sensory adaptation.

**Significance statement:** Vertebrate vision depends on opsins, which detect specific wavelengths of light. Most teleosts retain a single long-wavelength–sensitive opsin (LWS1), while a green-sensitive paralog (LWS2) occurs only in a few lineages. We identify a previously overlooked green-shifted opsin, LWS3, which is restricted to gobies, and show that LWS2 and LWS3 have repeatedly evolved green sensitivity through convergent amino acid changes. In parallel to the primate MWS/LWS evolution, these shorter-shifted opsins have functionally replaced the canonical green-sensitive RH2 opsin in multiple teleost lineages. By integrating genomic, structural, and expression data, we reveal how multilevel convergent evolution—from molecular tuning to photoreceptor specialization—has repeatedly shaped green-light vision over the past 500 million years of evolution, illustrating the remarkable flexibility of vertebrate visual systems.

## Introduction

Animals rely on vision for a variety of fundamental tasks, including foraging, mating, avoiding predators, and navigating their environment. At the molecular level, the visual process is initiated when light is absorbed by photopigments, located in the outer segments of the retinal photoreceptors (Wald 1968; Hunt et al. 2014). Vertebrate photopigments consist of light-sensitive vitamin-A-derived chromophores, which are covalently bound to one of five different visual opsin protein types. These five types were already present in the vertebrate ancestor, and are thought to be the product of two rounds of whole-genome duplication (2R) ∼ 500 million years ago (Mya) (Lamb, 2013). In the retina, they are expressed in an ancestral set of photoreceptor types (PR0 – PR4) (Baden et al. 2025). They are the dim-light active rod opsin (rhodopsin; RH1, expressed in PR0) and four cone-based photopigments [SWS1 (PR4), SWS2 (PR3), RH2 (PR2), and MWS/LWS (PR1)], which are active during bright light and used for colour and motion vision (Hunt et al. 2014). Their spectral sensitivity ranges are a remarkable example of niche partitioning, where each cone-opsin pigment occupies a constricted visual space, spanning from the ultraviolet (<400 nm; SWS1) to the red end of the light spectrum (>600 nm; LWS) (Marshall et al. 2024).

The evolution of vertebrate visual opsins is characterised by changes in opsin gene numbers, gene structure and gene expression (Hunt et al. 2014; Lamb 2019). Unlike in terrestrial vertebrates, where the basic opsin setup was either maintained [e.g., birds (Borges et al., 2015), reptiles (Katti et al., 2019)] or reduced [e.g., mammals (Jacobs, 2013)], opsin genes have continued to proliferate in teleost fishes (Musilova et al., 2021; Policarpo et al. 2025). With over 35,000 species, teleosts are not only the most species-rich clade of vertebrates (Fricke et al., 2021), but they also exhibit extreme ecological, physiological, and morphological diversity, having conquered almost every aquatic and some terrestrial habitats (Cortesi et al., 2018). It is these differences in photic environment and life history traits that are the primary drivers of opsin gene diversity and evolution in fishes (Carleton et al. 2020).

Opsin gene evolution affects teleost vision at all taxonomic levels, from species to families and entire classes (Cortesi et al. 2015; Lin et al. 2017; Musilova et al. 2019; reviewed in Carleton et al. 2020; Musilova et al., 2021). Novel opsins commonly arise through tandem duplication, but, like the ancestral five types, may also be the product of whole-genome duplications (Musilova et al. 2021; Policarpo et al. 2025). Moreover, in some rare cases, they arise from retrotransposition (Mano et al. 1999; Ward et al. 2008). While most genes are lost shortly after duplication and disappear or become non-functional pseudogenes (Walsh, 2003), some genes persist by acquiring novel functions. This neofunctionalization of visual opsins typically involves changes in ‘key’ amino acid sites in the retinal binding pocket, which shift the spectral sensitivity of the photopigment and are often associated with the colonisation of novel light habitats or visual niches (Yokoyama 2008; Carleton et al. 2020). Neofunctionalization can also occur due to differences in gene expression with ontogeny (Carleton et al. 2008; Cortesi et al. 2016), location in the retina or other tissues (Davies et al. 2015; Morrow et al. 2016), or by changing non-visual characteristics of the protein (e.g. Castiglione et al., 2018; Luk et al., 2016). Finally, teleost opsins are prone to gene conversion, a form of reticulate evolution whereby sequence information is unidirectionally exchanged between sister genes (Cortesi et al. 2015; Sandkam et al. 2017). Gene conversion has the effect of homogenising paralogs, which may repair and sometimes even resurrect gene function (Cortesi et al. 2015).

The genomes of extant teleosts contain a median of seven visual opsin genes: one rod opsin and six cone opsins (Musilova et al. 2019). Most of the cone opsin expansions can be attributed to more recent lineage or species-specific tandem duplications of the *rh2* gene, which is sensitive to the most common, blue to green light spectrum underwater (peak spectral sensitivity range, λ_max_ ∼ 450 to 540 nm) (Carleton et al. 2020; Musilova & Cortesi, 2023). Several older tandem duplications have also persisted, such as those found in the neoteleost and percomorph violet to blue-sensitive *sws2* duplicates (Cortesi et al. 2015). Interestingly, at their origin ∼ 320 Mya, teleosts underwent a third round of whole-genome duplication (3R) (Ravi and Venkatesh 2018), and two extant opsin gene duplicates, one *rh1* (Chen et al., 2018) and one *lws* (Liu et al. 2019), have been attributed to this event. The ancestral *lws*2 duplicate has long been overlooked but has recently been described from bony tongues (Osteglossiformes) and characins (Characiformes) (Liu et al. 2019; Escobar-Camacho et al. 2020).

Most teleosts possess a single LWS visual pigment (LWS1), which is sensitive from the yellow to red spectrum (λ_max_ ∼ 560 to 620 nm) (Carleton et al. 2020; Schweikert et al., 2018), or they might have lost the gene altogether, as commonly seen in deep-sea lineages (Musilova et al. 2019; Lupše et al., 2021). The second copy, LWS2, in extant species, but likely already in the ancestor shortly after its duplication by 3R, has shifted its spectral sensitivity to shorter, greener wavelengths (λ_max_ ∼ 530 nm) (Escobar-Camacho et al. 2020). This is similar to the MWS/LWS duplicate in primates, which re-enabled trichromatic vision after the loss of *rh2* in the mammalian ancestor (Carvalho et al., 2017). Concurrently, both bony tongues and characins have either lost or severely reduced the expression of *rh2*, suggesting that there is a trade-off between the overlap of pigment spectral sensitivities and the potential for evolutionary retention (Liu et al. 2019; Escobar-Camacho et al. 2020). Interestingly, a shift towards shorter spectral sensitivities can also be observed in some more recent LWS1 duplicates in several fish species, as well as between different alleles of the same gene (Seehausen et al. 2008; Sandkam et al. 2018). Given the repeated pattern of *lws* duplication followed by occupation of the green-light niche, it is surprising that teleosts only possess a limited number of these genes. Moreover, the convergence of green-sensitive LWS visual pigments in distantly related vertebrate groups presents a unique opportunity to examine how evolution is repeated to achieve similar visual functions.

Against this background, we explore the evolutionary history of the *lws* opsins in teleost fishes from a phylogenetic representative sampling of more than 100 species. By combining next-generation genome and transcriptome sequencing with open-source data mining, we uncover a rich history of gene duplication, loss and gene conversion. Among these, we find two more teleost groups: tarpons (Elopomorpha), which also retain the *lws2* duplicate, and gobies (Percomorpha), which have a third *lws* copy (*lws3*) of ancestral origin. We then focus on the (re)evolution of green sensitivity and investigate the protein-level signatures of parallelism between the teleost green-sensitive LWS2 and LWS3 visual pigments and the human MWS, uncovering remarkable signatures of repeated amino acid substitutions and functional convergence in gene expression patterns across the vertebrate tree of life.

## Results and Discussion

To reconstruct the evolutionary history of LWS opsins, we analyzed available whole-genome data and sequenced retina transcriptomes from 20 selected species carrying a second, short-wavelength–shifted LWS paralog, primarily gobies and osteoglossiforms (Table S1). Combined with structural protein modeling, these data allowed us to focus on the (re)evolution of green sensitivity and the convergent emergence of green–red LWS opsin pairs in teleost fishes and humans.

### *Lws* gene duplications and synteny

Reconstructing a comprehensive *lws* gene tree based on 109 teleost genomes and adding non-teleost ray-finned (Actinopterygian), lobe-finned (Sarcopterygii), and several ancestrally derived fishes as outgroups, we confirm that most teleosts have one red-sensitive *lws* gene (*lws1*). In addition, we find that the tarpons (Elopomorpha) have retained the *lws2* copy, which was previously reported only from bony tongues and characins, and emerged during 3R (Liu et al. 2019; Escobar-Camacho et al. 2020). Surprisingly, however, we uncover a third ancestral *lws* (*lws3*) gene in gobies (Gobiiformes) (Figs. 1, 2, and S1). The teleost-specific *lws* duplication that gave rise to *lws2* is currently the earliest described cone opsin duplication (reviewed in Musilova et al., 2021); our results show that *lws3* is possibly of a similar or even more ancient origin. However, accurately dating the *lws3* duplication is more challenging than for *lws2*, mainly due to insufficient phylogenetic signal in the data. Three scenarios are likely:

1. *Lws3* is an ancient duplicate predating the teleost *lws1–lws2* duplication, which is supported by the nucleotide tree (Fig. S1). However, this finding may be influenced by the saturation of the third-codon positions (Breinholt & Kawahara, 2013), making the phylogenetic signal questionable. Moreover, as gobies are a crown-percomorph group, such an ancestral origin would require at least 19–20 independent *lws3* losses across teleosts (Fig. 1).
2. *Lws3* is homologous to *lws2* (i.e., derived from the same ancestral duplicate); however, there is no phylogenetic support for this scenario besides for exon 3, which groups the goby *lws3* together with the exon 3 of *lws2* (Fig. S4). This hypothesis also lacks any syntenic support (Fig. S3), and, similar to scenario 1, it implies the independent loss of *lws2/lws3* in all non-goby euteleosts. Nevertheless, five identical amino-acid pairs in the retinal-binding pocket of the LWS2 and LWS3 proteins (Fig. 2B) might suggest otherwise, albeit these might have arisen convergently.
3. *Lws3* is a more recent duplication originating in the percomorph ancestor. Supporting this scenario, to recover the monophylum with goby *lws1* and *lws3* always involves other percomorphs based on amino-acid sequences or reduced nucleotide trees (excluding the third codon position), although these trees remain largely unresolved (Fig. S5). It would also suggest an unusually rapid sequence evolution in goby *lws3* and convergence with *lws2*, which is not found in extant goby species.

**Figure 1:**
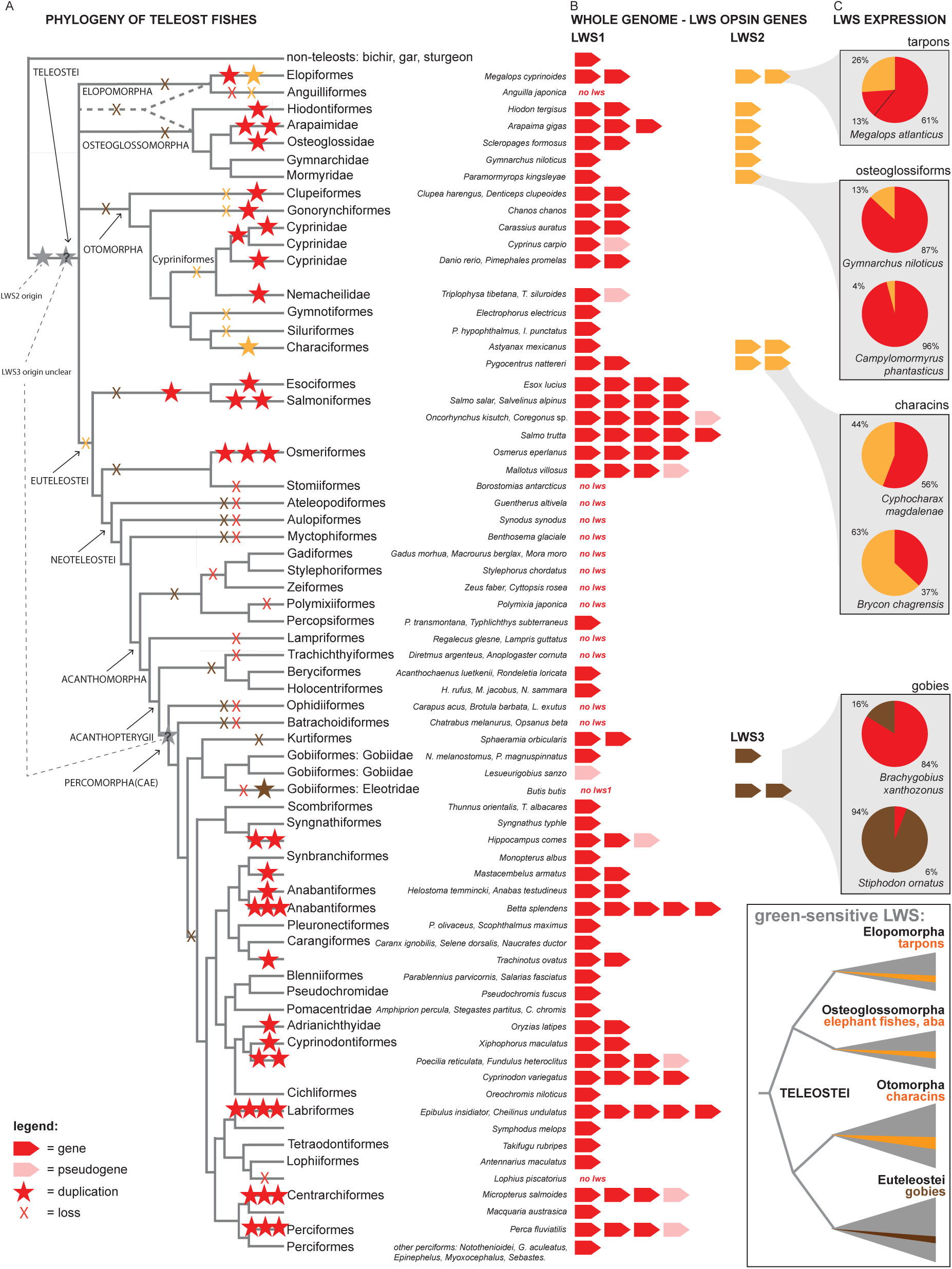
*Lws* opsin gene repertoire in teleost fishes. A) Putative gene duplications (star symbols) and gene losses (X symbols) mapped onto the teleost phylogeny (simplified after Betancur et al., 2017). Most duplications are species/genus specific, except for in the Esociformes and Salmoniformes, and the Cyprinodontiformes, which are older. Two ancient gene duplications are marked at the onset of teleosts (grey stars). B) The number and type of *lws* genes found in the whole genomes of teleost fishes. Red - *lws1* (red-sensitive), orange - *lws2*, brown - *lws3* (both shifted towards shorter, green wavelengths). Note that only a few lineages, including Elopomorpha, Osteoglossomorpha and Characiformes, retain *lws2*, and only gobies (Gobiiformes) have *lws3*. C) Proportional gene expression in representative species with multiple *lws* paralogs: tarpon (Elopomorpha), elephant fishes (Osteoglossiformes), characins (Ostariophysi), and gobies (Percomorpha). The position of these groups within the teleost phylogeny is highlighted in the inset. Refer to Fig. S1 for the *lws* gene tree, Table S1 for the list of analysed genomes, and Fig. 5 for the details on the goby *lws3* evolutionary history.

**Figure 2:**
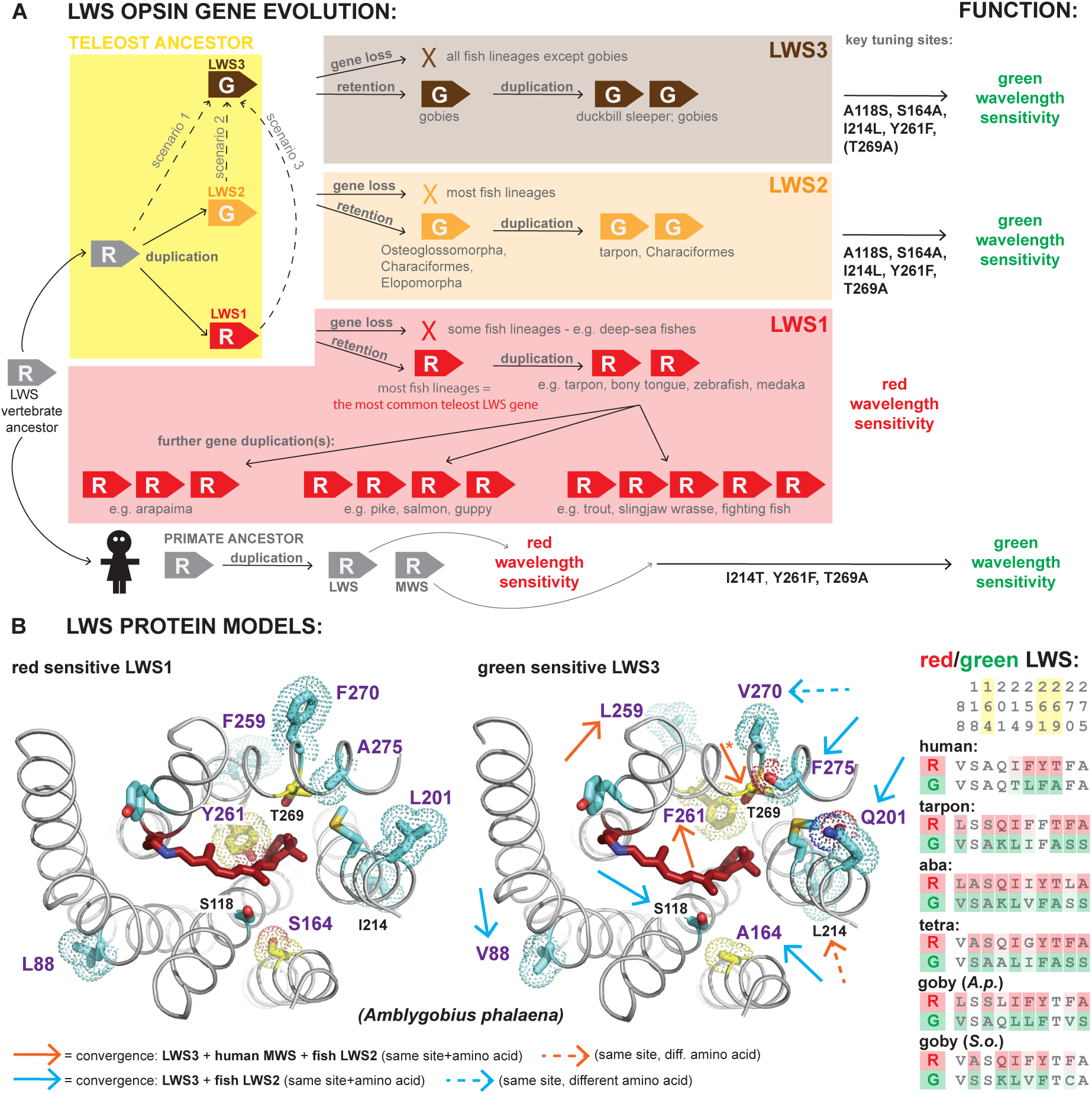
Convergent evolution of green-light-sensitive LWS opsins. A) Proposed model of the *lws* gene dynamics in teleost fishes. *Lws1* is present in most fish lineages and is commonly sensitive to longer (red) wavelengths >560 nm. The *lws2* copy is retained in Osteoglossomorpha, Elopomorpha and Characiformes (and is absent from Euteleostei). *Lws3* occurs in gobies (Gobiiformes) and has presumably been lost in other fish lineages. Both LWS2 and LWS3 are functionally shifted towards shorter, green wavelengths ∼520 nm. B) LWS opsin protein models by Boltz (Passaro et al. 2025). Across vertebrates, green sensitivity evolved convergently from a red-sensitive ancestor via changes in key amino acids. Depicted are three amino acid sites that show the same mutations in human MWS (green) opsin and in both teleost LWS2 and LWS3 opsins (red arrows). Further sites have evolved convergently among the teleost LWS2 and LWS3 paralogs (blue arrows). * marks the mutation T269A, which is found in humans, teleost LWS2, and in LWS3 of mudskippers, but not in the two goby species selected for the modelling. For all 12 structural models performed in this study, please refer to Fig. S8.

An additional and unexpected outcome of our phylogenetic analyses is that *lws* genes from some tetrapods (amphibians, snakes, lizards, and mammals) and the non-tetrapod lungfish cluster within the teleost *lws* clade, whereas *lws* genes from other tetrapods (birds and turtles) follow the species tree and branch outside the ray-finned fishes (Fig. S1). This pattern suggests the possibility of yet another ancestral *lws* duplication, potentially in the common ancestor of ray-finned and lobe-finned fishes. Under this scenario, birds and turtles would have retained a different *lws* paralog than other tetrapods. However, more extensive sampling across tetrapod lineages will be required to fully resolve this aspect of *lws* evolutionary history.

A detailed analysis of the synteny (±10 flanking genes) around teleost *lws* loci revealed a highly conserved genomic context for *lws1*, which is consistently located between *hcfc1* or *sws2* upstream and *gnl3l* downstream (Fig. S3). In contrast, *lws2* shows reduced syntenic conservation (Fig. 3) and occurs on distinct chromosomal regions in bony tongues and characins. Notably, we detected interspecific synteny between two *lws2* genomic regions in tarpon (Elopomorpha) and the *lws2* regions of *Paramormyrops* (Osteoglossomorpha) and cavefish (Characiformes). One tarpon *lws2* region closely matches the *Paramormyrops* cluster, whereas the second corresponds more closely to the characiform cluster (Figs. 3, S2, and S3). This pattern implies that the teleost ancestor possessed two *lws2* paralogs located in distinct genomic regions, both of which are retained in tarpons, while only one copy persists in osteoglossomorphs and the other in ostariophysans.

**Figure 3:**
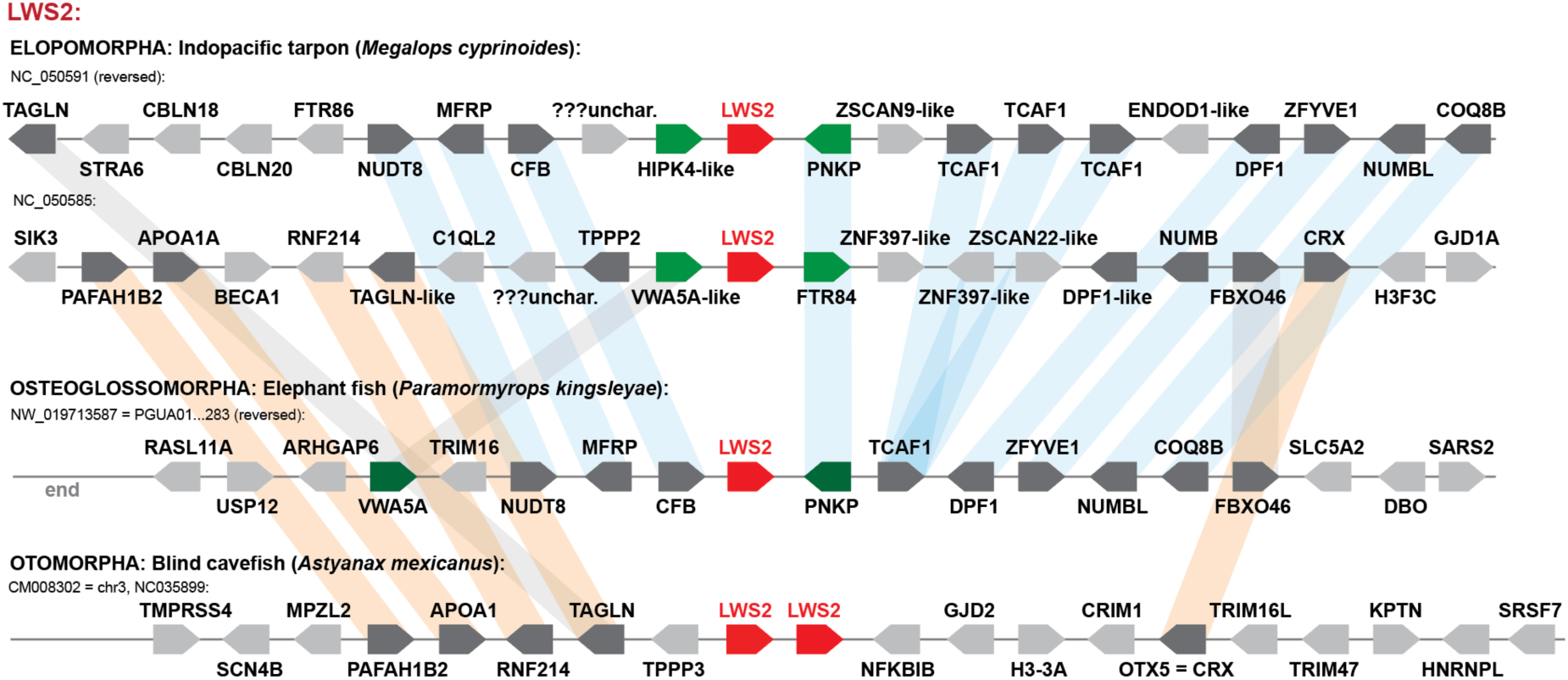
*Lws2* gene synteny. Two clusters with the *lws2* gene located on two different chromosomes are found in the Indopacific tarpon (Elopomorpha). One cluster is present in each of the elephant fish (Osteoglossomorpha) and the blind cavefish (Characiformes, Otomoropha). One of the tarpon clusters (NC_050591) shares more syntenic features with the elephant fish cluster (genes connected by blue shading). The second cluster (NC_050585) is more like the cavefish cluster (connected by orange bars). Note that the *lws2* genes themselves no longer carry this phylogenetic signal (see gene trees in Figs. 4 and S1). For the synteny of a longer genomic fragment (up to 2 Mbp), refer to Fig. S2. See Fig. S3 for the synteny of *lws1* and *lws3*.

In gobies, *lws3* is located in a distinct genomic region, flanked by *rab32* upstream and *stxbp5* or *androglobin* downstream (Fig. S3). This region lacks any opsin gene in other teleosts (Fig. S3), and no conserved synteny is detectable among the *lws1*, *lws2*, and *lws3* loci. One possibility is that both *lws2* and *lws3* have undergone genomic relocation over evolutionary time. Transposable elements are known to play a significant role in opsin gene dynamics in teleosts, as demonstrated in African cichlids (Carleton et al., 2020b), and both teleost *rh1* and one *lws1* copy in guppies and other livebearers originated via retrotransposition (Mano et al., 1999; Ward et al., 2008; Sandkam et al., 2017). Alternatively, the ancestral synteny may have been obscured by lineage-specific gene losses or extensive chromosomal rearrangements over deep evolutionary time.

A close inspection of the *lws* cis-regulatory elements revealed that the canonical vertebrate *lws1* transcription factor binding sites (OTX, THR, RAX; Ghinia Tegla et al., 2020; Emerson et al., 2013; Volkov et al., 2020; Mackin et al., 2019; Nelson et al., 2009; Furukawa et al., 1997) are absent from the upstream regions of *lws2* and *lws3* (Table S2 and Fig. S7). Moreover, putative miRNA signals detected upstream of one *lws2* copy in *Megalops* and *lws3* in *Neogobius* exhibit low folding energies (miRNAfold; Tav et al., 2016) and limited similarity to vertebrate miR726, indicating distinct regulatory architectures for these paralogs, whose functional elements remain to be identified (Table S2 and Fig. S7).

### The dynamics of the *lws* evolutionary history

Most shallow marine and freshwater fishes retain at least one copy of *lws1* (Figure 1), whereas many deep-living taxa, including anglerfishes, cusk-eels, toadfishes, and the deep-water goby *Lesueurigobius sanzi*, have lost *lws* genes entirely. This pattern is consistent with the rapid attenuation of long wavelengths with depth, which likely renders *lws* functionally obsolete in deep-water environments. In contrast, *lws1* duplications are common in lineages inhabiting red-shifted or visually complex environments [e.g. some characins (Escobar-Camacho et al. 2020)] or using orange and yellow colouration, such as wrasses (Phillips et al., 2016) and livebearers (Windsor & Owens, 2009). Multiple paralogs are also arising through lineage-specific whole-genome duplications (e.g., Cypriniformes, Salmoniformes, Esociformes) or tandem duplications and reaching up to five *lws1* paralogs in species such as brown trout, Siamese fighting fish, and several wrasses (Figure 1).

Gene conversion between *lws* paralogs has been documented for *lws1* copies in guppies and between *lws1* and *lws2* in characins (Escobar-Camacho et al., 2020; Sandkam et al., 2017). Similarly, we found evidence for gene conversion in exon 3 of the *lws1* and *lws2* opsins in tarpons (Fig. S4). Furthermore, GARD (Kosakovsky Pond et al., 2006) revealed conversion between *lws1* and *lws3* in the peacock gudgeon *Tateurndina ocellicauda* (Fig. S6), resulting in two genes with highly similar functional properties (Fig. 5). However, for the remaining gobies, patterns of gene conversion were less pronounced than those reported for other opsin duplicates, such as *sws2* (Cortesi et al., 2015). This suggests that *lws3* diverged rapidly after duplication, with subsequent gene conversion acting as a partial homogenizing force—primarily affecting the 5′ portion of the coding sequence—while exons containing spectral tuning sites remained comparatively unaffected.

**Table 1:**
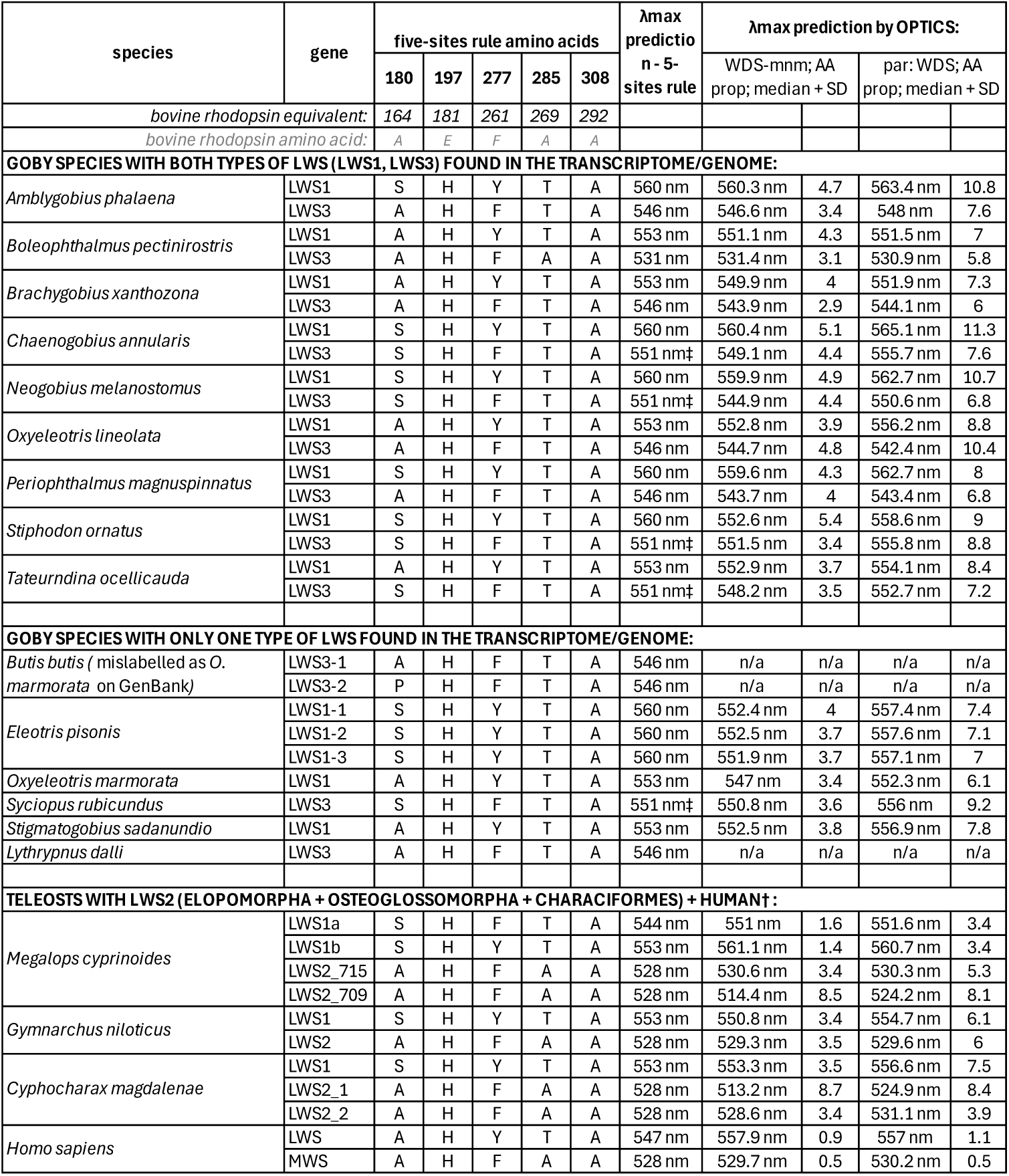
Predicted LWS peak spectral sensitivities (λ_max_) in gobies (LWS1, LWS3) and other teleosts (groups with LWS2). For the gobies, we started with the predicted value (λ_max_ = 551 nm) for a mudskipper LWS opsin (You et al. 2014), based on the five-site rule with S164-H181-F261-T269-A292 (marked with ‡). We added +6 nm in 164, +9 nm in 261, and +13 nm in 269 when a polar residue was present (Sekharan et al., 2012; Yokoyama, 2008). For species with LWS2 (*Megalops*, characin, *Gymnarchus*), we started with the value from bovine rhodopsin ∼ 500 nm (+28 nm for E181H). OPTICS (Frazer and Oakley, 2025) was used for the machine-learning predictions on the entire dataset (mine-and-match or heterologous), considering amino-acid properties (AA_prop parameter). Generally, the sensitivity of LWS2 and LWS3 is shifted towards green compared to LWS1. The bovine rhodopsin sites 164, 181, 261, 269, and 292 correspond to sites 180, 197, 277, 285, and 308 in the human LWS (Hagen et al., 2023).

### Convergent red to green functional shifts in vertebrate LWS visual pigments

Similar to the repeated shift from red- to green-sensitivity in mammals (Carvalho et al., 2017; Melin et al., 2016), we find that shorter-wavelength-shifted LWS opsins have evolved repeatedly in teleost fishes (Table 1). Importantly, amino acid sites involved in this shift, including those for the human MWS-LWS pair, have undergone convergent changes across vertebrates (Hagen et al., 2023). In teleosts, this has previously been described for several derived teleost LWS1 copies and LWS2 in characins and bony tongues (Liu et al., 2019; Escobar-Camacho et al., 2020; Sandkam et al., 2018). Here, we describe a convergent shorter-wavelength shift in LWS2 of the tarpons and the goby LWS3 (Fig. 2A; Table 1).

The molecular evolution of these copies was assessed using structural models for one opsin pair from each of six species: human (*Homo sapiens*, MWS and LWS), the osteoglossomorph aba (*Gymnarchus niloticus*, LWS1 and LWS2), the elopomorph tarpon (*M. cyprinoides*, LWS1 and LWS2), the ostariophysan characin (*Cyphocharax magdalenae*, LWS1 and LWS2), and two percomorph gobies (*Amblygobius phalaena* and *Stiphodon ornatus*, both LWS1 and LWS3). The models focused on amino acid variation within the retinal-binding pocket, encompassing ten variable sites (Figs. 2B and S8; Table 2).

**Table 2:** Variable position in the retinal binding pocket. The five-site rule amino acid positions, 164, 261, and 269, are highlighted in yellow. Predictions for peak spectral sensitivity (λ_max_) values using the five-site rule are provided.

| bovine rhodopsin position | $\lambda_{\max}$ / | 88 | 118 | 164 | 201 | 208 | 214 | 259 | 261 | 269 | 270 | 275 | 293 |
| --- | --- | --- | --- | --- | --- | --- | --- | --- | --- | --- | --- | --- | --- |
| human_NM_020061_LWS_OPN1LW | 547 | V | S | A | Q | M | I | F | Y | T | F | A | Y |
| human_NM_000513_MWS_OPN1MW | 528 | V | S | A | Q | M | T | L | F | A | F | A | F |
| Aphalaena_FZ1_LWSx1 | 560 | L | S | S | L | M | I | F | Y | T | F | A | Y |
| Aphalaena_FZ3_LWSx2 | 546 | V | S | A | Q | M | L | L | F | T | V | S | Y |
| Stiphodon_ornatus_LWS01 | 560 | V | A | S | Q | M | I | F | Y | T | F | A | Y |
| Stiphodon_ornatus_LWS02 | 551 | V | S | S | K | M | L | V | F | T | C | A | Y |
| Cmagdalenae_MT310731 | 553 | V | A | S | Q | M | I | G | Y | T | F | A | Y |
| Cmagdalenae_MT310746 | 528 | V | S | A | A | L | L | I | F | A | S | S | Y |
| Gymnarchus_niloticus_119C1_LWS1 | 553 | L | A | S | Q | M | I | I | Y | T | L | A | Y |
| Gymnarchus_niloticus_119C1_LWS2 | 528 | V | S | A | K | M | L | V | F | A | S | S | Y |
| Megalops_cyprinoides_CM023712_LWS01A | 544 | L | S | S | Q | M | I | F | F | T | F | A | Y |
| Megalops_cyprinoides_CM023709_LWS2 | 528 | V | S | A | K | L | L | I | F | A | S | S | Y |

We identified three key positions: Y261F, T269A, and F259L (bovine rhodopsin numbering) that show clear signatures of convergence between human and teleost LWS opsins; in addition, site 214 consistently distinguishes green- and red-sensitive opsins, albeit via different amino acid substitutions (Fig. 2B). Sites 261 and 269 are well-established contributors to vertebrate LWS spectral tuning and form part of the “five-sites rule” or the “OH-sites rule” (Yokoyama and Radlwimmer, 1998; Yokoyama, 2008; Sekharan et al., 2012; Chi et al., 2020). In addition, an extra five amino acid substitutions (L88V, A118S, Q201K, 270 variable, and A275S), and changes at site 164 (member of the five-sites rule), were common between green-shifted LWS2 and LWS3 opsins (Fig. 2B). These shared substitutions support the convergent evolution between the two teleost LWS paralogs (unless *lws2* and *lws3* are homologous; see scenario 2, above).

Despite convergently shorter shifting of the newly emerged LWS copies, ancestral-state reconstruction of the three most variable tuning sites (164, 261, and 269) revealed functional divergence among the paralogs (Fig. 4). LWS1 consistently retains red sensitivity by mainly preserving longer-wavelength–shifted residues, whereas LWS2 predominantly carries green-shifted variants at these three sites. LWS3 exhibits an intermediate pattern: site 261 is green-shifted in all gobies, site 164 is variably green-shifted among species, and site 269 is green-shifted only in mudskippers, with other gobies retaining the longer-wavelength variant (Fig. 4; Table 1). This was supported by a machine-learning approach (OPTICS; Frazer and Oakley, 2025) to predict the spectral sensitivities of the LWS opsins from 15 goby species, as well as from three teleost species with LWS2 (Table 1). Estimated λ_max_ values for goby LWS1 ranged from 551.9 to 567.3 nm, whereas green-shifted LWS2 ranged from 517 to 529 nm, and LWS3 was intermediate, ranging from 523 to 557 nm. These predicted values largely overlapped with measured photoreceptor sensitivities reported for other goby species, including the shorter-shifted double cone with λ_max_ = 527–531 nm (Pierotti et al., 2020, and references therein), which is thought to express LWS3 only.

**Figure 4:**
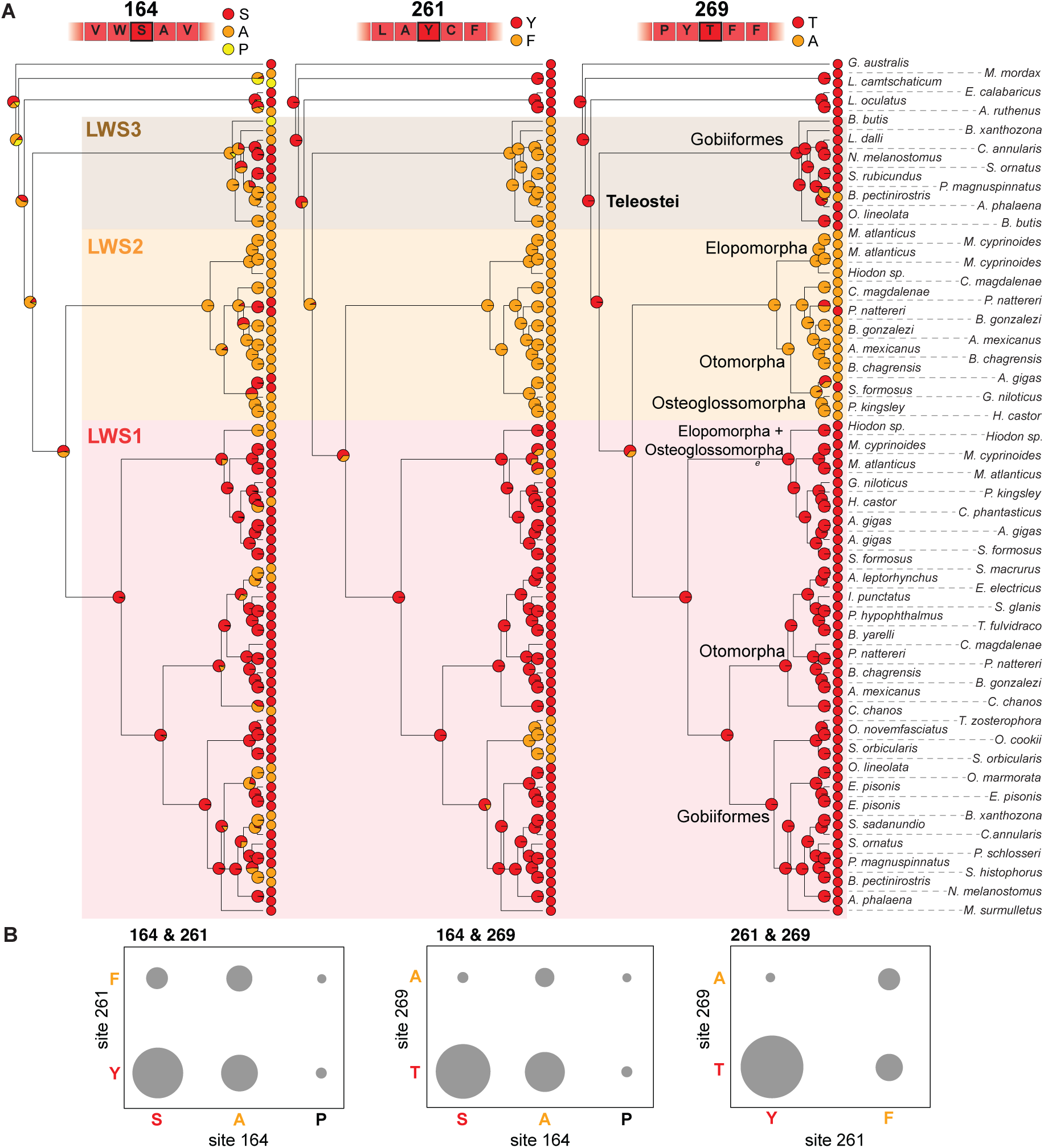
Ancestral state reconstruction of the three most variable key-tuning sites of the teleost LWS paralogs. A) Sites 164, 261 and 269 are known to have a large effect on LWS spectral sensitivity (Musilova et al., 2021). LWS1 has mostly longer-shifted amino acid variants at all three sites (red circles). LWS2 has shorter-shifted amino acid variants (orange circles), resulting in a notable shift towards green spectral sensitivity. LWS3 lies in between, with a shorter-shifted variant (F) at site 261, a predominantly longer-shifted variant (T) at site 269, and a combination of both variants at site 164 (S and A) or an even shorter variant (P) encoded in yellow. Pie charts at the nodes indicate the scaled likelihoods of each variant, calculated using the ace function in APE (Paradis & Schliep, 2019). B) Association of amino acid variants in the teleost LWS evolution (325 haplotypes). Most copies are long-wavelength-sensitive (red) at the majority of sites, reflecting the dominance of LWS1 in teleosts.

### Neofunctionalization of LWS3 in gobies

We confirmed the functional relevance of multiple LWS copies across teleost lineages by examining their retinal expression patterns. Consistent with their predicted distinct spectral sensitivities, *lws1* and *lws2* were found to be expressed in the retinas of the tarpon (*Megalops*; Elopomorpha), gymnarchids (Gymnarchidae) and elephantfishes (Mormyridae; both Osteoglossomorpha), and in characins (Characiformes; Otomorpha) (Fig. 1C). Similarly, both *lws1* and *lws3* were expressed in gobies (Gobiiformes), although the expression patterns varied markedly between species (Figs. 1C and 5).

**Figure 5:**
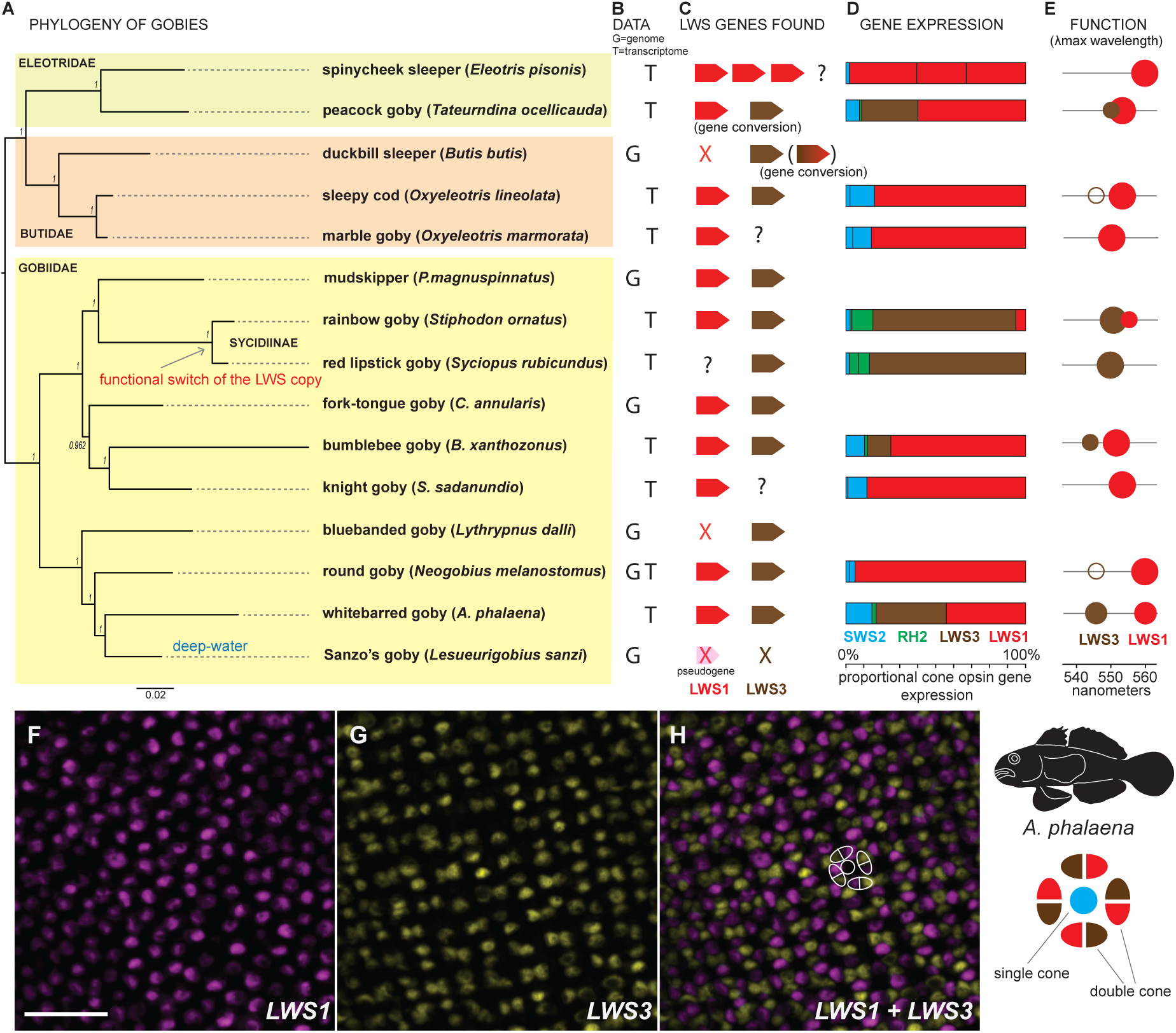
*Lws* evolution in gobies (Gobiiformes). A) Phylogenetic relationship between studied gobies based on the genomic and transcriptomic data set (three families, n = 15 species). B) Data source: genomic (G), transcriptomic (T), both (GT). C) Mined *lws* genes, where each polygon represents one copy (*lws1*, red polygon; *lws3*, brown polygon) and an X marks gene loss. Note that no conclusion about gene losses can be drawn based on transcriptomic data alone (marked by ?). D) Proportional cone opsin gene expression in the gobies. The colour code in the horizontal bars represents the spectral sensitivity range of the corresponding visual pigment: blue = *sws2*, green = *rh2*, red = *lws1*, and brown = *lws3*. Note that either *lws1* (most gobies) or *lws3* (Sycidiinae) dominate the cone opsin gene expression profiles. E) The spectral sensitivity (λ_max_) of the LWS1 (red) and LWS3 (brown) opsins based on the predictions in Table 1. Larger versus smaller circles represent the dominant versus rare transcripts, and an empty circle represents no expression, based on data in D). Note the functional proximity of LWS1 and LWS3 in the peacock goby (*Tateurndina ocellicauda*), in which gene conversion occurred (see also Fig. S6). F-H) Double-labelling fluorescent *in situ* hybridization of expressed opsin mRNAs in retinal double cone photoreceptors in adult *Amblygobius phalaena. Lws1* (magenta) and *lws3* (yellow) mRNAs were expressed in opposite members of double cones—scale bars: 10 µm. For the *in situ* of the single cones, refer to Fig. S9.

Comparing the retinal transcriptomes of ten goby species representing nine genera and most major subfamilies revealed that *lws1* was the most highly expressed paralog in most taxa, with five species expressing *lws3* to varying degrees (Fig. 5). An exception was found in the subfamily Sycidiinae: the rainbow goby (*Stiphodon ornatus*) predominantly and the red lipstick goby (*Syciopus rubicundus*) exclusively expressed *lws3*, respectively (Fig. 5C). In addition to *lws*, all gobies expressed at least one copy of *sws2*, albeit at low levels in the Sycidiinae and in the spinycheek sleeper (*Eleotris pisonis*). In contrast, the green-sensitive opsin RH2 was generally absent or expressed only at trace levels, except for the Sycidiinae, where *rh2* expression was comparable to *sws2* levels in other gobies (Fig. 5C). Taken together with the modeled green sensitivity of LWS3 (Fig. 2; Table 1), the reduced reliance on *rh2* supports a functional replacement of RH2-mediated green vision by LWS3 in gobies, providing strong support for the neofunctionalization of this paralog.

Neofunctionalization of *lws3* was further supported in *A. phalaena*, where, using fluorescent in situ hybridization, we found that *lws1* and *lws3* are expressed exclusively in opposite members of the double cones (Fig. 5 F–H; Fig. S9 D–F). On the other hand, the *sws2* opsins (*sws2b* and *sws2aβ*) were expressed exclusively in single cones (Fig. S9 A, B). *Sws2aβ* was expressed in nearly all photoreceptors, whereas *sws2b* showed variable and often lower expression, resulting in partial coexpression within the same single cones (Fig. S9 A–C). This cellular segregation demonstrates that *lws3* has acquired an independent photoreceptor role in *A. phalaena* and, together with *sws2*- and *lws1*-expressing photoreceptors, likely underpins trichromatic vision in this species.

## Conclusion

By reconstructing the evolutionary history of *lws* opsins across ray-finned fishes, we show that repeated gene duplication, divergence, and differential retention have generated three ancient *lws* paralogs—*lws1*, *lws2*, and *lws3*—that remain functional in extant lineages despite extensive genomic rearrangement and regulatory divergence. Our discovery of *lws3* reveals an unrecognized layer of sensory diversity and demonstrates that ancient duplicates can be repeatedly co-opted to expand visual function long after their origin. In gobies, LWS3 has undergone neofunctionalization to mediate green sensitivity, functionally replacing the canonical green-sensitive opsin RH2. This role parallels that of LWS2 in osteoglossiforms and characins and mirrors primate color vision, in which the duplication of *lws* enabled green-sensitive cones following the loss of *rh2* in the mammalian ancestor. Across these systems, convergent amino acid substitutions at key spectral-tuning sites underpin parallel functional shifts, highlighting common molecular solutions to similar selective pressures. Together, our results show that opsin gene evolution in teleost fishes is exceptionally dynamic, with ancient gene duplicates serving as a recurrent substrate for sensory innovation. This evolutionary flexibility contrasts with the more constrained opsin repertoires of other vertebrates, underscoring the power of comparative genomic and functional approaches to reveal hidden aspects of visual system evolution.

## Materials and Methods

### Ethical approval and relevant permits

All experimental procedures involving live animals within Australia were approved by the University of Queensland Animal Ethics Committee (QBI/304/16). Fish were collected under the Great Barrier Reef Marine Parks Permit (G17/38160.1) and Queensland general fisheries permits (207976 and 199434). Sample collection in the Czech Republic was conducted under a permit issued by the Czech Fisheries Association. Specimens collected for retinal transcriptomics were euthanized using MS222 or an ice slurry, in accordance with best practice standards.

### Taxon sampling

Samples of *Amblygobius phalaena* and *Epibulus insidiator* for gene expression analysis have been previously published in Musilova et al., 2019 (see Table S1 for the sample list and Accession numbers). *Oxyeleotris lineolata* (n = 4 sub-adults) were collected from the Dawson River at Taroom, Queensland, Australia, in May 2020. *Amblygobius phalaena* specimens (n = 3 adults) for in-situ hybridisation were collected on snorkel from shallow reefs (1-2 m) surrounding Lizard Island, Great Barrier Reef, Australia, in March 2021 using an anaesthetic clove oil solution (10% clove oil, 40% ethanol, and 50% seawater) and hand nets. Eyes were enucleated, and the retinas were dissected out and either fixed in 4% PFA for *in situ* hybridization or stored in RNAlater at −20 °C until further processing for transcriptome sequencing. Samples of *Neogobius melanostomus* were collected in the Elbe River (Děčín, Czech Republic) using an electrofishing device. Samples of the remaining seven species, *Eleotris pisonis, Eleotris marmorata, Tateurndina ocellicauda, Stiphodon ornatus, Syciopus rubicundus, Brachygobius xanthozonus* and *Stigmatogobius sadanundio*, were obtained from the aquarium trade.

#### Transcriptome sequencing

Retinal RNA was extracted using a Qiagen mini kit column extraction following the manufacturer’s protocol and including a DNAse step. RNA extractions concentrations were measured on a Fluorometer and integrity checked with a Pico chip on a 2100 Bioanalyzer from Agilent. Library preparation (unstranded, 300 bp insert) was conducted using Illumina’s NEBNext Ultra II Directional RNA library preparation kit, NEBNext Multiplex Oligos and the NEBNext Poly(A) mRNA Magnetic Isolation Module (New England Biolabs) for all osteoglossiform, tarpon and goby species used for transcriptomics except for *Oxyleotris lineolata, Neogobius melanostomus* and *Amblygobius phalaena* whose library preparations have been performed on the commercial basis in Novogene, Singapore (https://en.novogene.com/). Transcriptome sequencing (RNAseq HiSeq Paired-End PE150) was outsourced to Novogene for all species.

#### Assembly and gene extraction

Transcriptome raw reads have been filtered and mapped with Medium sensitivity settings against the general goby reference (based on the genome of *Neogobius melanostomus*; Adrian-Kalchauser et al., 2019) in the Geneious 9.0 software (https://www.geneious.com). Species specific gene sequences have been retrieved from the mapped alignments and used as a species-specific reference for another round of mapping with the Lowest sensitivity settings. Number of reads have been calculated to the FPKM (fragments per kilo base per million reads) and the proportion of each opsin from the total cone opsin gene expression has been calculated. More details on the protocols described in de Busserolles et al. 2017, Musilova et al., 2019, and Tettamanti et al. 2019.

#### Whole genome analysis

High quality teleost whole genomes have been downloaded from GenBank (accession numbers as per sample Table S1). Majority of the high-quality genomes has been sequenced by the Sanger Institute (Rhie et al., 2021). We further complemented this data set by the genomic data of lower quality (draft genomes as per Cortesi et al., 2015 and Musilova et al., 2019). Assembled genomic scaffolds / chromosomes have been mapped against the general single exon reference of the example LWS1 (*Oreochromis niloticus*), LWS2 (*Astyanax mexicanus*) and LWS3 (*Neogobius melanostomus*) copy. Mapping has been performed in Geneious 9.1.8 (https://www.geneious.com) using Medium to Medium High sensitivity settings - this settings has been tested to pick all scaffolds with any LWS opsin (higher sensitivity yielded to false hits). Single LWS genes have been extracted from the scaffolds either following the annotation (if available) or by mapping of single exons against the scaffold with the Highest sensitivity.

#### Phylogenetic analysis

Coding sequences of the single LWS genes have been aligned in Geneious 9.1.8 using the MAFFT Alignment plugin (Katoh et al., 2019). Five independent runs (each with two parallel runs) have been performed in MrBayes 3.1.2 (Ronquist and Huelsenbeck, 2003) software for 50 million generations. The output files have been manually inspected in the Tracer 1.5 (https://github.com/beast-dev/tracer/releases) for reaching the lnL plateau, the best -lnL score and to identify the burnin break point. Three out of the ten independent output runs have been used for the consensus tree construction with the burnin set to 50%.

#### Ancestral state reconstruction

In order to assess the evolution of the key-tuning amino acid sites, we focused on the three key sites that are known to cause large shifts in λ_max_ (sites 164, 261 and 269 – 7,10 and 16 nm, respectively) and reconstructed the ancestral states for each key site with the gene tree pruned to the taxa with multiple LWS types, i.e., the Elopomorpha+Osteoglossomorpha+Otomorpha with LWS1 and LWS2 and for GObiiformes with LWS1 and LWS3 gene types. Using the occurrence of amino acids variants that shift λ_max_ as character states, and with the equal transition rates model (ER), we performed the ancestral state analysis the ace function in the APE package (Paradis & Schliep, 2019). The ace function employs a maximum likelihood approach where the reconstructed ancestral states are given as a proportion of the total likelihood for each state at each node. Trees were also analysed, visualized, and edited with the phytools package.

#### Gene conversion analysis

GARD (Kosakovsky Pond et al. 2006) was run using general discrete site-to-site variation with three rate classes on the Datamonkey platform (www.datamonkey.org) (Weaver et al. 2018) on an alignment of *LWS1* and *LWS3* from goby species for which both copies were recovered (Fig. 3). The sliding window analysis followed the methods described in Cortesi et al. 2015. In short, the intraspecific dS (neutral process) along the coding sequences of *LWS1* and *LWS3* was calculated using a window size of 30 and a step size of 1 in DNAsp v.6 (Rozas et al. 2017).

#### Regulatory elements analysis

The key regulatory elements controlling expression of the *LWS* opsin gene were examined including the locus control region (LCR), which is shared across vertebrates (Smallwood et al 1992; Wakefield et al 2018), and the associated miRNA (miR-726) shared across fishes (O’Quin et al 2011; Smallwood et al., 2002; Wakefield et al., 2008). In most fish, the miRNA and LCR are within a 500 bp region. We aligned these regions from *Oryzias latipes, Gasterosteus aculeatus, Oreochromis niloticus* and *Danio rerio,* to identify the core LCR and miRNA regions. These were then blasted to the genomes of *Astyanax, Paramormyrops, Meglalops,* and *Neogobius* to identify homologous regions upstream of any of the *LWS* genes. Within the LCRs, we identified binding sites for key regulators of the LWS gene (OTX2, THR or RAX; Ghinia Tegla et al., 2020; Emerson et al., 2013; Volkov et al., 2020; Mackin et al., 2019; Nelson et al., 2009; Furukawa et al., 1997) using Jaspar 2020 (Fornes et al., 2019). If there was homologous sequence to miR-726, we tested miRNA stability by calculating the miRNA folding energy with miRNAfold (https://evryrna.ibisc.univ-evry.fr/miRNAFold). In this method, a more negative energy signifies more stability and so a more likely functional miRNA.

#### Protein structure prediction and retinal binding pocket analysis

Three-dimensional structures of LWS opsins were predicted using Boltz (Passaro et al., 2025), a deep learning-based protein structure prediction method. Predicted structures were converted from mmCIF to PDB format for downstream analysis. Bovine rhodopsin (PDB: 1GZM, chain A) and human OPN1LW (NM_020061) and OPN1MW (NM_000513) were included as structural references.

The retinal chromophore binding pocket was characterized by identifying residues within 6 Å of 11-cis-retinal and its covalently bound lysine. Residues were categorized into three distance shells from the retinal-lysine complex: close (<4 Å), medium (4–5 Å), and far (5–6 Å). Analysis was performed using PyMOL (Schrödinger, LLC) with custom Python scripts. Opsin sequences were aligned to establish positional homology across species. Binding pocket residues identified by distance criteria were mapped to alignment positions, enabling cross-species comparison of equivalent sites. Residues at positions corresponding to the "five-sites rule" for LWS/MWS spectral tuning (positions 180, 197, 277, 285, and 308 in human opsin numbering) were extracted from the binding pocket analysis.

#### In-situ hybridisation

Dual-labelling RNA-FISH was performed on two wholemount retinas (left and right) of one adult *A. phalaena* collected at Lizard Island on 16 March 2021 following standard protocols (e.g., Barthel and Raymond 2000, Raymond and Barthel 2004, Allison et al. 2010, Dalton et al. 2014, Dalton et al. 2015). Retinas were prepared as follows. After one-hour dark adaptation, the eyes were enucleated, vitreous was enzymatically digested, and retinas were extracted from the eyecups. The retinal pigment epithelium layer (RPE) was removed by PBS-jetting using a single use syringe and needle (Terumo). Retinas were flattened and pinned, photoreceptor side down, in Sylgard-filled petri dishes, and fixed in 4% PFA at 4°C overnight. Fixed retinas were triple washed in PBS for 10 min, briefly rinsed in 70% methanol in PBS, and then transferred to 100% methanol and stored at −20°C.

Previously extracted retinal mRNA was reverse transcribed using a High Capacity RNA-to-cDNA Reverse Transcription Kit (Applied Biosystems). Riboprobe templates were synthesized from cDNA via standard PCR using opsin specific primers (Table S3) and Q5 High Fidelity DNA polymerase (New England Biolabs). Primers were designed to bind to the coding sequence of target opsins (*SWS2B*, *SWS2Aβ*, *LWS1*, *LWS3*) and to contain a T3 or T7 RNA polymerase promoter sequence at their 5’-ends (T3, reverse primer; T7, forward primer). Amplicons were isolated via gel-electrophoresis and gel-extraction (Qiagen Gel Extraction Kit), and subsequently used as template in enrichment PCR using the same primers. Anti-sense riboprobes labelled with either digoxigenin-UTP (DIG) or fluorescein-UTP (FL) were synthesized from cDNA templates via T3 RNA polymerase mediated strand-specific RNA transcription using DIG/FL RNA labelling mix (Roche). Hybridised, labelled riboprobes were detected using anti-digoxigenin or anti-fluorescein antibodies conjugated to horseradish peroxidase (Roche). Fluorescent tagging was performed using Alexa Fluor 594 and 488 dyes with streptavidin Tyramide Signal Amplification (TSA, Invitrogen). Finally, retinas were mounted in 70% glycerol in PBS, photoreceptor side up, on microscopy slides and covered with a coverslip.

Fluorescent tagged photoreceptor cells were imaged using a CFI Apo Lambda 60x/1.4 NA oil immersion objective on a spinning disk confocal microscope (Diskovery, Andor Technologies, UK) built around a Nikon Ti-E body (Nikon Corporation, Japan) equipped with two Zyla 4.2 sCMOS cameras (Andor Technology), and controlled by Nikon NIS Elements software (Nikon Corporation, Japan). Exported images were further processed (max. intensity z-stack projection, channel merging) with ImageJ v. 1.52p (National Institute of Health, USA).

## Data Availability

All sequenced transcritpomic data have been submitted to GenBank and are available under accession numbers listed in the Table S1.

## Acknowledgements

We would like to thank Dr Andrew Mather for help with sample collection. We thank Veronika Truhlářová for lab management and logistical support. FC was supported by the Australian Research Council DECRA (DE200100620) and Future (FT240100725) Fellowships and by a University of Queensland Development Fellowship, Australia. GMS has been funded by GAUK (1524119; Charles University) and SVV 260-792/2025. ZM was supported by the Swiss National Science Foundation (SNF grant PROMYS, 166550), Charles University (Primus), Czech Science Foundation (21-31712S) and an ERC Consolidator grant (SensingDeep).

## Author contributions

Z.M. and F.C. designed research and analyzed data; D.E.C. performed the ancestral state reconstruction, C.K. performed the tests of regulatory regions and contributed to the overall interpretations. G.M.S. performed the RNA extractions and transcriptome sequencing. F.H. and X.D. performed the structural protein modelling. M.L. performed the fluorescent in situ hybridization. Z.M. and F.C. wrote the initial version of the manuscript. All authors have contributed to the advanced versions of the manuscript.

## References

Adrian-Kalchhauser, I., Blomberg, A., Larsson, T., Musilova, Z., Peart, C. R., Pippel, M.,… Winkler, S. (2020). The round goby genome provides insights into mechanisms that may facilitate biological invasions. BMC Biology, 18(1), 1–33. 10.1186/s12915-019-0731-8

Baden, T., Angueyra, J. M., Bosten, J. M., Collin, S. P., Conway, B. R., Cortesi, F.,… & Corbo, J. C. (2025). A standardized nomenclature for the rods and cones of the vertebrate retina. PLoS Biology, 23(5), e3003157.

Borges, R., Khan, I., Johnson, W. E., Gilbert, M. T. P., Zhang, G., Jarvis, E. D.,… & Antunes, A. (2015). Gene loss, adaptive evolution and the co-evolution of plumage coloration genes with opsins in birds. BMC genomics, 16(1), 1–14.

Breinholt, J. W., & Kawahara, A. Y. (2013). Phylotranscriptomics: saturated third codon positions radically influence the estimation of trees based on next-gen data. Genome Biology and Evolution, 5(11), 2082–2092.

Castiglione, G. M., Schott, R. K., Hauser, F. E., & Chang, B. S. (2018). Convergent selection pressures drive the evolution of rhodopsin kinetics at high altitudes via nonparallel mechanisms. Evolution, 72(1), 170–186.

Carleton, K.L., Conte, M.A., Malinsky, M., Nandamuri, S.P., Sandkam, B.A., Meier, J.I., Mwaiko, S., Seehausen, O. and Kocher, T.D., 2020. Movement of transposable elements contributes to cichlid diversity. Molecular Ecology, 29(24), pp.4956–4969.

Carleton, K.L., Escobar-Camacho, D., Stieb, S.M., Cortesi, F. and Marshall, N.J., 2020. Seeing the rainbow: mechanisms underlying spectral sensitivity in teleost fishes. Journal of Experimental Biology, 223(8).

Carvalho, L. S., Pessoa, D., Mountford, J. K., Davies, W. I., & Hunt, D. M. (2017). The genetic and evolutionary drives behind primate color vision. Frontiers in Ecology and Evolution, 5, 34.

Chen J-N, Samadi S, Chen W-J. 2018. Rhodopsin gene evolution in early teleost fishes. PLOS ONE. 13(11):e0206918

Chi, H., Cui, Y., Rossiter, S.J. and Liu, Y., 2020. Convergent spectral shifts to blue-green vision in mammals extends the known sensitivity of vertebrate M/LWS pigments. Proceedings of the National Academy of Sciences, 117(15), pp.8303–8305.

Cortesi, F., Cheney, K. L., Cooke, G. M., & Ord, T. J. (2018). Opsin gene evolution in amphibious and terrestrial combtooth blennies (Blenniidae). BioRxiv, 503516.

Cortesi, F., Musilová, Z., Stieb, S.M., Hart, N.S., Siebeck, U.E., Malmstrøm, M., Tørresen, O.K., Jentoft, S., Cheney, K.L., Marshall, N.J. and Carleton, K.L., 2015. Ancestral duplications and highly dynamic opsin gene evolution in percomorph fishes. Proceedings of the National Academy of Sciences, 112(5), pp.1493–1498.

Cortesi, F., Musilová, Z., Stieb, S.M., Hart, N.S., Siebeck, U.E., Cheney, K.L., Salzburger, W. and Marshall, N.J., 2016. From crypsis to mimicry: changes in colour and the configuration of the visual system during ontogenetic habitat transitions in a coral reef fish. Journal of Experimental Biology, 219(16), pp.2545–2558.

Davies, W. I., Tamai, T. K., Zheng, L., Fu, J. K., Rihel, J., Foster, R. G.,… & Hankins, M. W. (2015). An extended family of novel vertebrate photopigments is widely expressed and displays a diversity of function. Genome research, 25(11), 1666–1679.

de Busserolles, F., Cortesi, F., Helvik, J.V., Davies, W.I., Templin, R.M., Sullivan, R.K., Michell, C.T., Mountford, J.K., Collin, S.P., Irigoien, X. and Kaartvedt, S., 2017. Pushing the limits of photoreception in twilight conditions: the rod-like cone retina of the deep-sea pearlsides. Science advances, 3(11), p.eaao4709.

Emerson, M. M., Surzenko, N., Goetz, J. J., Trimarchi, J., & Cepko, C. L. (2013). Otx2 and Onecut1 promote the fates of cone photoreceptors and horizontal cells and repress rod photoreceptors. Developmental cell, 26(1), 59–72.

Escobar-Camacho, D., Carleton, K.L., Narain, D.W. and Pierotti, M.E., 2020. Visual pigment evolution in Characiformes: the dynamic interplay of teleost whole-genome duplication, surviving opsins and spectral tuning. Molecular Ecology, 29(12), pp.2234–2253.

Fornes O, Castro-Mondragon JA, Khan A, et al. JASPAR 2020: update of the open-access database of transcription factor binding profiles. Nucleic Acids Res. 2019; doi: 10.1093/nar/gkz1001

Frazer, S. A., & Oakley, T. H. (2025). Accessible and Robust Machine Learning Approaches to Improve the Opsin Genotype-Phenotype Map. bioRxiv, 2025-08.

Fricke, R., Eschmeyer, W. N. & Van der Laan, R. (eds) 2021. ESCHMEYER’S CATALOG OF FISHES: GENERA, SPECIES, REFERENCES. (http://researcharchive.calacademy.org/research/ichthyology/catalog/fishcatmain.asp). Electronic version accessed 02/04/2021.

Furukawa, T., Kozak, C. A., & Cepko, C. L. (1997). Rax, a novel paired-type homeobox gene, shows expression in the anterior neural fold and developing retina. Proceedings of the National Academy of Sciences, 94(7), 3088–3093.

Galbraith, J.D., Ludington, A.J., Suh, A., Sanders, K.L. and Adelson, D.L., 2020. New Environment, New Invaders—Repeated Horizontal Transfer of LINEs to Sea Snakes. Genome Biology and Evolution, 12(12), pp.2370–2383.

Ghinia Tegla, M. G., Buenaventura, D. F., Kim, D. Y., Thakurdin, C., Gonzalez, K. C., & Emerson, M. M. (2020). OTX2 represses sister cell fate choices in the developing retina to promote photoreceptor specification. elife, 9, e54279.

Hagen, J. F., Roberts, N. S., & Johnston Jr, R. J. (2023). The evolutionary history and spectral tuning of vertebrate visual opsins. Developmental Biology, 493, 40–66.

Hunt DM, Hankins MW, Collin SP, Marshall NJ. 2014. Evolution of Visual and Non-Visual Pigments. Boston: Springer

Jacobs, G. H. (2013). Losses of functional opsin genes, short-wavelength cone photopigments, and color vision--a significant trend in the evolution of mammalian vision. Visual Neuroscience, 30(1-2), 39.

Jerlov NG. 1976. Marine Optics. Amsterdam: Elsevier. 2nd ed.

Katoh, K., Rozewicki, J., & Yamada, K. D. (2019). MAFFT online service: multiple sequence alignment, interactive sequence choice and visualization. Briefings in bioinformatics, 20(4), 1160–1166.

Katti, C., Stacey-Solis, M., Coronel-Rojas, N. A., & Davies, W. I. L. (2019). The diversity and adaptive evolution of visual photopigments in reptiles. Frontiers in Ecology and Evolution, 7, 352.

Kosakovsky Pond, S.L., Posada, D., Gravenor, M.B., Woelk, C.H. and Frost, S.D., 2006. Automated phylogenetic detection of recombination using a genetic algorithm. Molecular biology and evolution, 23(10), pp.1891–1901.

Lamb, T.D., 2013. Evolution of phototransduction, vertebrate photoreceptors and retina. Progress in retinal and eye research, 36, pp.52–119.

Lamb, T.D., 2019. Evolution of the genes mediating phototransduction in rod and cone photoreceptors. Progress in retinal and eye research, p.100823.

Lin, J.J., Wang, F.Y., Li, W.H. and Wang, T.Y., 2017. The rises and falls of opsin genes in 59 ray-finned fish genomes and their implications for environmental adaptation. Scientific reports, 7(1), pp.1–13.

Liu, D.W., Wang, F.Y., Lin, J.J., Thompson, A., Lu, Y., Vo, D., Yan, H.Y. and Zakon, H., 2019. The cone opsin repertoire of osteoglossomorph fishes: gene loss in mormyrid electric fish and a long wavelength-sensitive cone opsin that survived 3R. Molecular biology and evolution, 36(3), pp.447–457.

Luk, H. L., Bhattacharyya, N., Montisci, F., Morrow, J. M., Melaccio, F., Wada, A.,… & Olivucci, M. (2016). Modulation of thermal noise and spectral sensitivity in Lake Baikal cottoid fish rhodopsins. Scientific Reports, 6(1), 38425.

Lupše, N., Cortesi, F., Freese, M., Marohn, L., Pohlmann, J. D., Wysujack, K.,… & Musilova, Z. (2021). Visual Gene Expression Reveals a cone-to-rod Developmental Progression in Deep-Sea Fishes. Molecular Biology and Evolution, 38(12), 5664–5677.

Lynch, M., & Conery, J. S. (2000). The evolutionary fate and consequences of duplicate genes. Science, 290(5494), 1151–1155.

Mackin, R. D., Frey, R. A., Gutierrez, C., Farre, A. A., Kawamura, S., Mitchell, D. M., & Stenkamp, D. L. (2019). Endocrine regulation of multichromatic color vision. Proceedings of the National Academy of Sciences of the United States of America, 116(34), 16882–16891. 10.1073/pnas.1904783116.

Mano H, Kojima D, Fukada Y. 1999. Exo-rhodopsin: a novel rhodopsin expressed in the zebrafish pineal gland. Mol. Brain Res. 73(1–2):110–18

Marshall, J., Cronin, T., Johnsen, S., Douglas, R., Hurlbert, A., Boddy, J., & Cortesi, F. (2024). Color in Nature. Princeton University Press. ISBN 978-0-691-25861-4

Melin, A. D., Wells, K., Moritz, G. L., Kistler, L., Orkin, J. D., Timm, R. M.,… & Dominy, N. J. (2016). Euarchontan opsin variation brings new focus to primate origins. Molecular Biology and Evolution, 33(4), 1029–1041.

Morrow, J.M., Lazic, S., Fox, M.D., Kuo, C., Schott, R.K., Gutierrez, E.D.A., Santini, F., Tropepe, V. and Chang, B.S., 2017. A second visual rhodopsin gene, rh1-2, is expressed in zebrafish photoreceptors and found in other ray-finned fishes. Journal of Experimental Biology, 220(2), pp.294–303.

Musilova, Z., Cortesi, F., Matschiner, M., Davies, W.I., Patel, J.S., Stieb, S.M., de Busserolles, F., Malmstrøm, M., Tørresen, O.K., Brown, C.J. and Mountford, J.K., 2019. Vision using multiple distinct rod opsins in deep-sea fishes. Science, 364(6440), pp.588–592.

Musilova, Z., & Cortesi, F. (2023). The evolution of the green-light-sensitive visual opsin genes (RH2) in teleost fishes. Vision Research, 206, 108204.

Musilova Z., Salzburger, W., Cortesi, F. (2021). The Visual Opsin Gene Repertoires of Teleost Fishes: Evolution, Ecology and Function. Annual Review of Cell and Developmental Biology.

Nelson, S. M., Park, L., & Stenkamp, D. L. (2009). Retinal homeobox 1 is required for retinal neurogenesis and photoreceptor differentiation in embryonic zebrafish. Developmental biology, 328(1), 24–39.

O’Quin, K. E., Smith, D., Naseer, Z., Schulte, J., Engel, S. D., Loh, Y. H. E.,… & Carleton, K. L. (2011). Divergence in cis-regulatory sequences surrounding the opsin gene arrays of African cichlid fishes. BMC evolutionary biology, 11(1), 120.

Paradis, E., & Schliep, K. (2019). ape 5.0: an environment for modern phylogenetics and evolutionary analyses in R. Bioinformatics, 35(3), 526–528.

Passaro, S., Corso, G., Wohlwend, J., Reveiz, M., Thaler, S., Somnath, V. R.,… & Barzilay, R. (2025). Boltz-2: Towards accurate and efficient binding affinity prediction. bioRxiv 2025. Doi: 10.1101/2025.06.14.659707

Phillips, G. A., Carleton, K. L., & Marshall, N. J. (2016). Multiple genetic mechanisms contribute to visual sensitivity variation in the Labridae. Molecular biology and evolution, 33(1), 201–215.

Pierotti, M.E., Wandycz, A., Wandycz, P., Rebelein, A., Corredor, V.H., Tashiro, J.H., Castillo, A., Wcislo, W.T., McMillan, W.O. and Loew, E.R., 2020. Aggressive mimicry in a coral reef fish: The prey’s view. Ecology and Evolution, 10(23), pp.12990–13010.

Policarpo, M., Fogg, L. G., Cortesi, F., & Salzburger, W. (2025). Evolution of the non-visual and visual opsin gene repertoire in ray-finned fishes. Genome Biology and Evolution, evaf129.

Rabosky, D.L., Chang, J., Title, P.O., Cowman, P.F., Sallan, L., Friedman, M., Kaschner, K., Garilao, C., Near, T.J., Coll, M. and Alfaro, M.E., 2018. An inverse latitudinal gradient in speciation rate for marine fishes. Nature, 559(7714), pp.392–395.

Ravi, V. and Venkatesh, B., 2018. The divergent genomes of teleosts. Annual review of animal biosciences, 6, pp.47–68.

Rhie, A., McCarthy, S. A., Fedrigo, O., Damas, J., Formenti, G., Koren, S.,… & Jarvis, E. D. (2021). Towards complete and error-free genome assemblies of all vertebrate species. Nature, 592(7856), 737–746.

Revell, L. J. (2012). phytools: an R package for phylogenetic comparative biology (and other things). Methods in ecology and evolution, (2), 217–223.

Ronquist, F. and J. P. Huelsenbeck. 2003. MRBAYES 3: Bayesian phylogenetic inference under mixed models. Bioinformatics 19:1572–1574.

Rozas, J., Ferrer-Mata, A., Sánchez-DelBarrio, J.C., Guirao-Rico, S., Librado, P., Ramos-Onsins, S.E. and Sánchez-Gracia, A., 2017. DnaSP 6: DNA sequence polymorphism analysis of large data sets. Molecular biology and evolution, 34(12), pp.3299–3302.

Sandkam, B.A., Joy, J.B., Watson, C.T. and Breden, F., 2017. Genomic environment impacts color vision evolution in a family with visually based sexual selection. Genome biology and evolution, 9(11), pp.3100–3107.

Sandkam, B., Dalton, B., Breden, F. and Carleton, K., 2018. Reviewing guppy color vision: Integrating the molecular and physiological variation in visual tuning of a classic system for sensory drive. Current zoology, 64(4), pp.535–545.

Seehausen, O., Terai, Y., Magalhaes, I.S., Carleton, K.L., Mrosso, H.D., Miyagi, R., Van Der Sluijs, I., Schneider, M.V., Maan, M.E., Tachida, H. and Imai, H., 2008. Speciation through sensory drive in cichlid fish. Nature, 455(7213), pp.620–626.

Sekharan, S., Katayama, K., Kandori, H., & Morokuma, K. (2012). Color vision:“OH-site” rule for seeing red and green. Journal of the American Chemical Society, 134(25), 10706–10712.

Smallwood, P. M., Wang, Y., & Nathans, J. (2002). Role of a locus control region in the mutually exclusive expression of human red and green cone pigment genes. Proceedings of the National Academy of Sciences, 99(2), 1008–1011.

Tav, C., Tempel, S., Poligny, L., & Tahi, F. (2016). miRNAFold: a web server for fast miRNA precursor prediction in genomes. Nucleic acids research, 44(W1), W181–W184.

Tettamanti, V., de Busserolles, F., Lecchini, D., Marshall, N.J. and Cortesi, F., 2019. Visual system development of the spotted unicornfish, Naso brevirostris (Acanthuridae). Journal of Experimental Biology, 222(24).

Volkov, L. I., Kim-Han, J. S., Saunders, L. M., Poria, D., Hughes, A. E., Kefalov, V. J.,… & Corbo, J. C. (2020). Thyroid hormone receptors mediate two distinct mechanisms of long-wavelength vision. Proceedings of the National Academy of Sciences, 117(26), 15262–15269.

Wakefield, M. J., Anderson, M., Chang, E., Wei, K. J., Kaul, R., Graves, J. A. M.,… & Deeb, S. S. (2008). Cone visual pigments of monotremes: filling the phylogenetic gap. Visual Neuroscience, 25(3), 257–264.

Wald, G., 1968. The molecular basis of visual excitation. Nature, 219(5156), pp.800–807.

Walsh, B. (2003). Population-genetic models of the fates of duplicate genes. In Origin and Evolution of New Gene Functions (pp. 279-294). Springer, Dordrecht.

Ward MN, Churcher AM, Dick KJ, Laver CR, Owens GL, et al. 2008. The molecular basis of color vision in colorful fish: Four Long Wave-Sensitive (LWS) opsins in guppies (Poecilia reticulata) are defined by amino acid substitutions at key functional sites. BMC Evol. Biol. 8(1):210

Weaver, S., Shank, S.D., Spielman, S.J., Li, M., Muse, S.V. and Kosakovsky Pond, S.L., 2018. Datamonkey 2.0: a modern web application for characterizing selective and other evolutionary processes. Molecular biology and evolution, 35(3), pp.773–777.

Windsor, D. J., & Owens, G. L. (2009). The opsin repertoire of Jenynsia onca: a new perspective on gene duplication and divergence in livebearers. BMC Research Notes, 2(1), 159.

Yokoyama, S., 2008. Evolution of dim-light and color vision pigments. Annu. Rev. Genomics Hum. Genet., 9, pp.259–282.

Yokoyama, S. and Radlwimmer, F.B., 1998. The" five-sites" rule and the evolution of red and green color vision in mammals. Molecular biology and evolution, 15(5), pp.560–567.

You, X., Bian, C., Zan, Q., Xu, X., Liu, X., Chen, J., Wang, J., Qiu, Y., Li, W., Zhang, X. and Sun, Y., 2014. Mudskipper genomes provide insights into the terrestrial adaptation of amphibious fishes. Nature communications, 5(1), pp.1–8.

