## Supplementary material for "Turning red to green repeatedly: ancient LWS duplications drive convergent visual tuning in teleost fishes": Suplementary Material

**Table S1.** Samples and accession numbers used in this study.

*(table provided in a separate file)*

**Table S2:** *The LWS regulatory elements found in the LWS1 genes but not much in the LWS2 and LWS3 upstream regions of the same species of cavefish (A. mexicanus), elephant fish (P. kingsleyae), tarpon (M. cyprinoides) and goby (N. melanostomus). The miRNA folding energy determined with* [https://evryrna.ibisc.univ-evry.fr/miRNAFold](https://evryrna.ibisc.univ-evry.fr/miRNAFold)*. More negative energy means more stable and so more likely functional miRNA. The LCR binding sites (OTX2, THR or RAX) identified with Jaspar. Grey marks the non-tested combinations (gene not present). For more details on miRNA detection, please check Figure S5:*

| | LWS1 | | LWS2 | | LWS2.II | | LWS3 | |
|---|---|---|---|---|---|---|---|---|
| | miR726 | LCR | miR726 | LCR | miR726 | LCR | miR726 | LCR |
| *Astyanax mexicanus* (Ostariophysi) | Yes E=-69 kcal/mol | Yes 2OTX 2THR | | | No | No | | |
| *Paramormyrops kingsleyae* (Osteoglossomorpha) | Yes E=-67 | Yes 2OTX, 1THR, 1RAX | No | No | | | | |
| *Megalops cyprinoides* (Elopomorpha) | Yes E=-61.3 | Yes 2OTX, 1THR | No | No | Yes E=-32 kcal/mol | No | | |
| *Neogobius melanostomus* (Percomorpha) | Yes E=-59.2 | Yes 2OTX | | | | | Yes E=-24 kcal/mol | No |

**Table S3.** Primers used for probe template synthesis for the fluorescent in situ hybridization in *Amblygobius phalaena*. T7 (forward primer) and T3 (reverse primer) RNA polymerase promoter sequences are shown in small letters.

| Target gene<br>Probe length (bp)<br>Probe similarity (gene, %) | Primer | Sequence |
|---|---|---|
| *SWS2B*<br>547<br>*SWS2Aβ*, 75.4% | SWS2B_F1<br>SWS2B_R2 | 5'-TAATACGACTCACTATAGGGACAGGCCTTTTCTACTCTTTGTCT-3'<br>5'-AATTAACCCTCACTAAAGGGGGACTCGTTGTTGTACTTGTTGTT-3' |
| *SWS2Aβ*<br>740<br>*SWS2B*, 75.4% | SWS2Aβ_F1<br>SWS2Aβ_R1 | 5'-TAATACGACTCACTATAGGGTGGGAGGTTTTATCAACGCTCTTA-3'<br>5'-AATTAACCCTCACTAAAGGGAAAGTTTCTCCACGGTTATTGACC-3' |
| *LWS1*<br>740<br>*LWS3*, 81.7% | LWS1_F1<br>LWS1_R1 | 5'-TAATACGACTCACTATAGGGGCTCTGGATGTGTTTTGTGGTTAT-3'<br>5'- AATTAACCCTCACTAAAGGGAAAGCAGGCAAAGAATGTGTAAGG-3' |
| *LWS3*<br>690<br>*LWS1*, 81.7% | LWS3_F2<br>LWS3_R2 | 5'-TAATACGACTCACTATAGGGATTTTCGAGGGCTACACTGTCTC-3'<br>5'-AATTAACCCTCACTAAAGGGTCGTCCACCTCTTTTCCAAACAG-3' |

**Figure S1** - Phylogenetic gene tree and key tuning amino acids of LWS. LWS2 and LWS3 genes are highlighted in orange and brown, respectively.

*(figure provided as a separate pdf file)*

**Figure S2**: Vista plot of the elephant fish (*P. kingsleyae*) and a cavefish (*A. mexicanus*) *LWS2* regions compared to two tarpon (*M. cyprinoides*) *LWS2* regions located on different chromosomes.

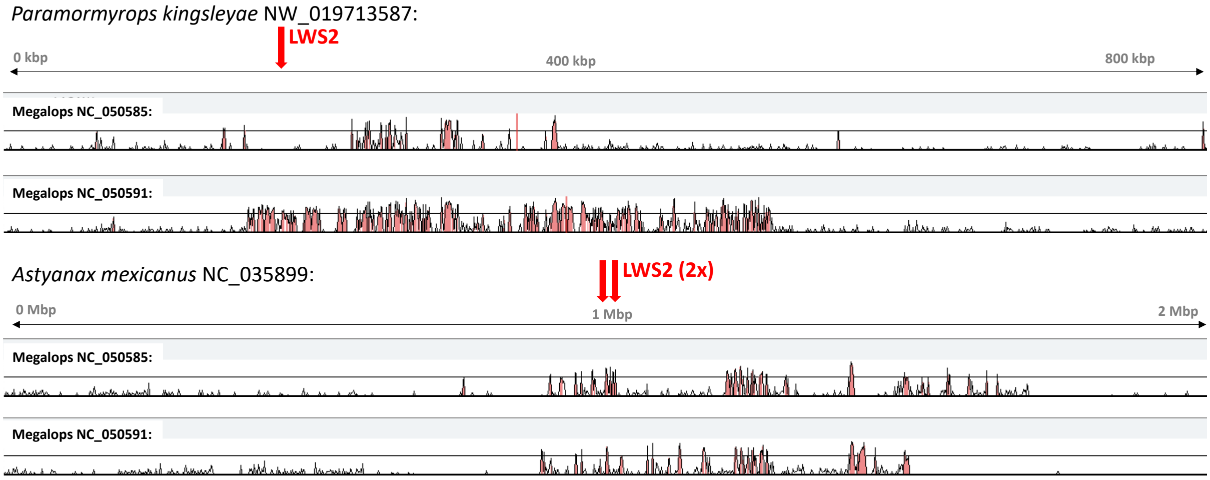

**Figure S3** - Synteny of the *LWS1*, *LWS2* and *LWS3* genes and the flanking regions in the round goby (*N. melanostomus*), elephant fish (*P. kingsleyae*), blind cavefish (*A. mexicanus*) and tarpon (*Megalops cyprinoides*).

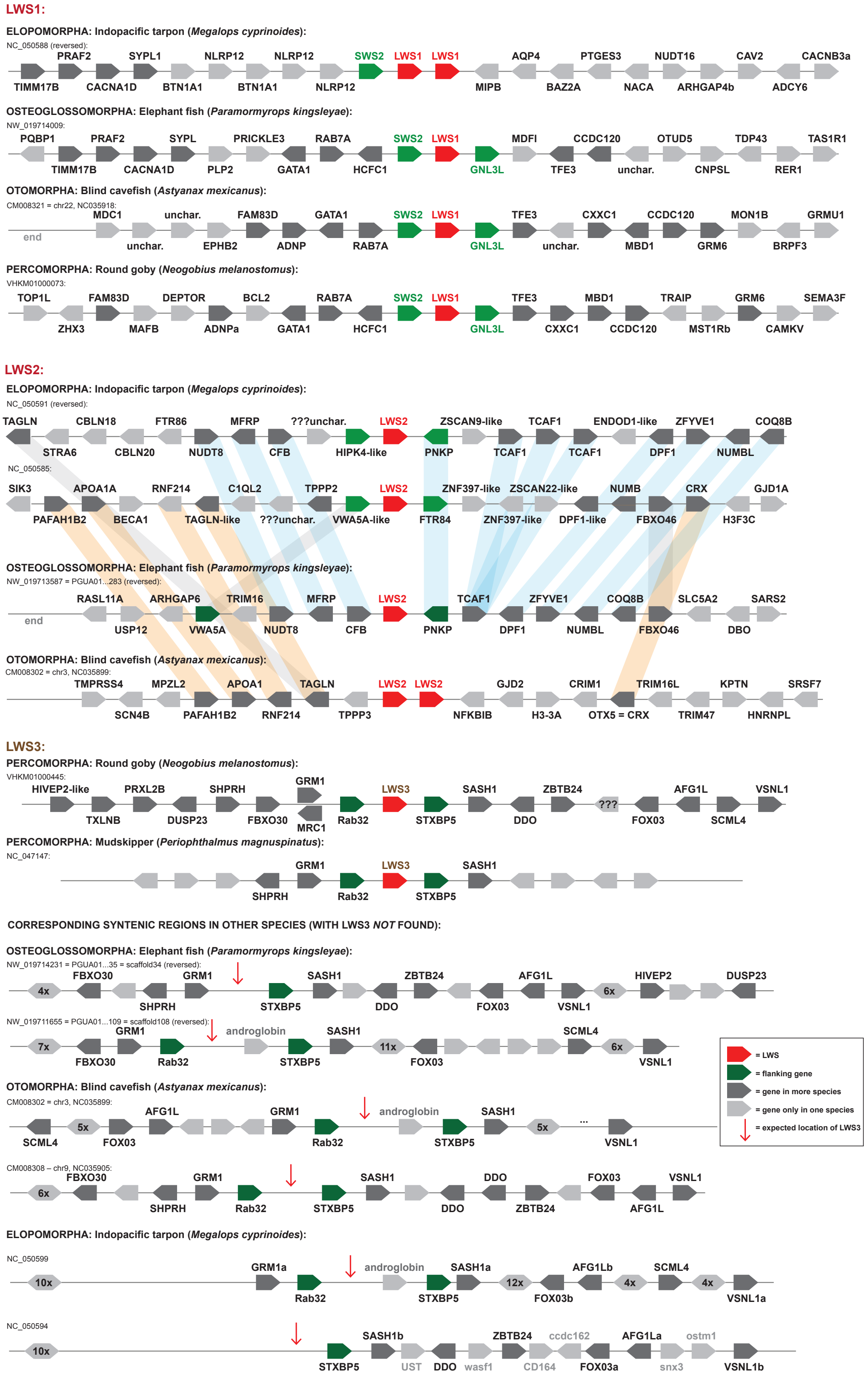

**Figure S4**: Single-exon gene tree (MrBayes). Orange highlights the *lws2* of osteoglossiform, characin and elopomorph fishes, blue marks goby *lws3* opsins. Note that in exon 3, *Megalops lws1* and *lws2* are similar, most likely due to gene conversion. In exon 4, the *lws3* and *lws2* form a grouping, although not corresponding to the species tree.

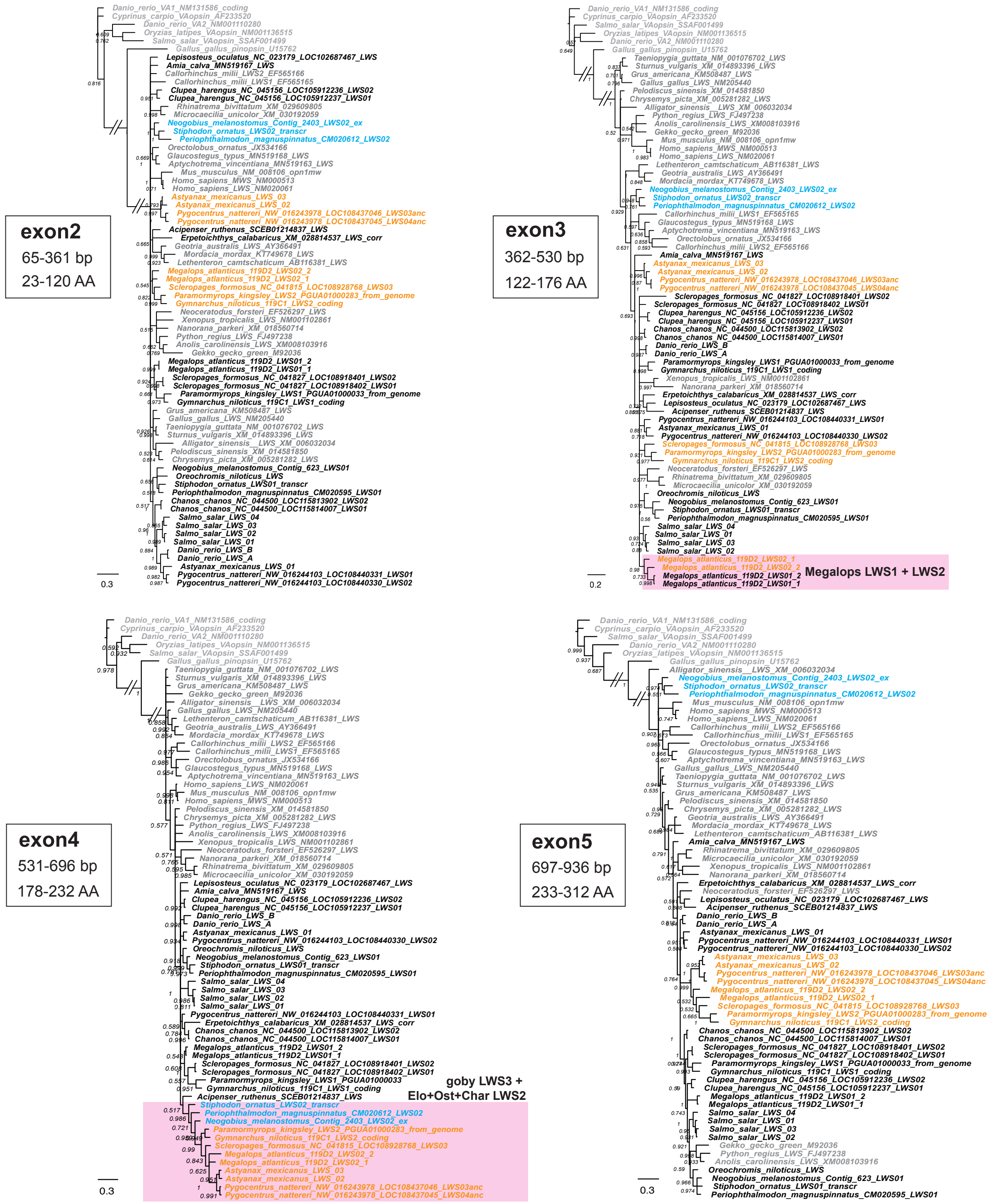

**Figure S5**: Phylogenetic analysis without the third codon position shows unresolved topology that involves percomorph fishes mixed with gobies.

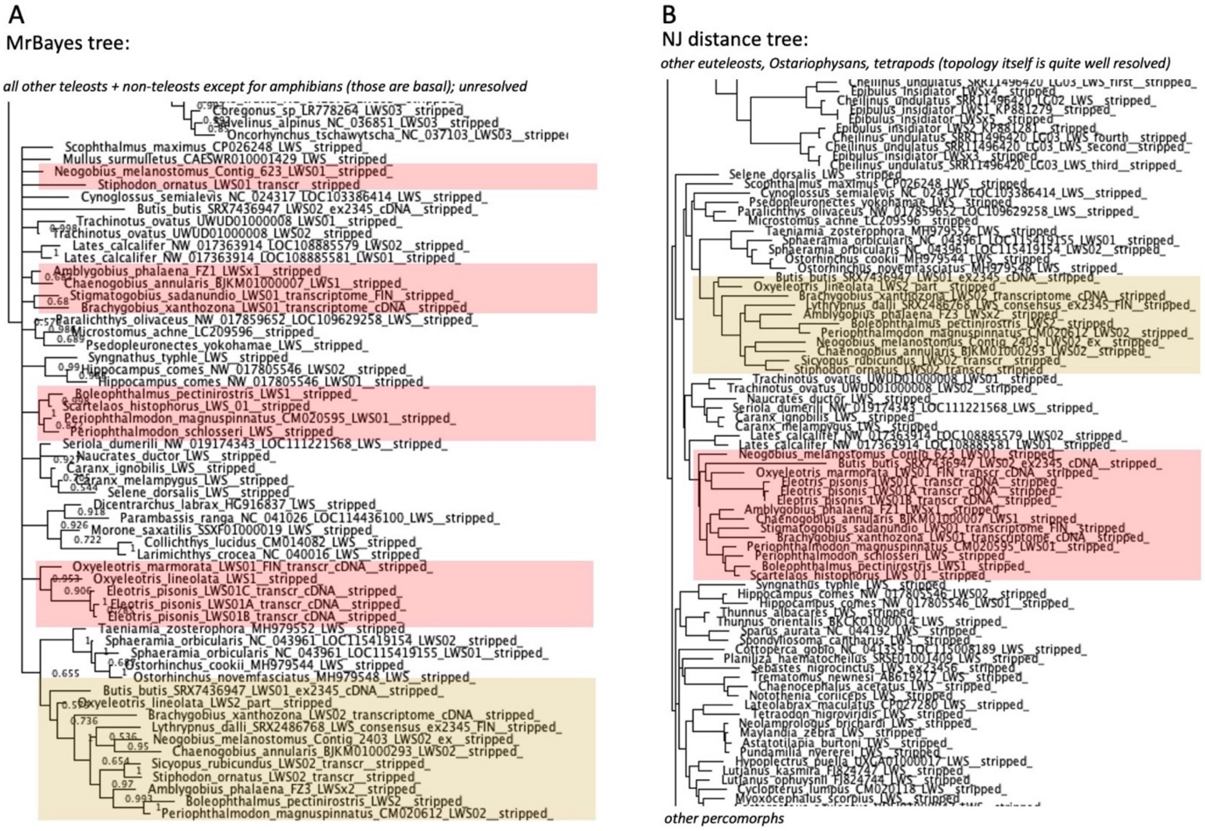

**Figure S6**: Gene conversion among the two *LWS* copies (*LWS1* and *LWS3*) of the goby, *Tateurndina ocellicauda*. Large-scale more ancient gene conversion impacted exons 2-3 and exons 5-6. These two sequences compromised the phylogenetic signal within the overall gene tree and had to be removed from the final analysis. A) Identity plot of the two nucleotide sequences. Green bars mark identical stretches of the coding region, white "gaps" indicate differences in sequence. Yellow boxes highlight the putative regions of gene conversion i.e., regions of high sequence similarity. B) GARD results for the comparison of all goby *LWS1* and *LWS3* sequences. The most likely breakpoints for *T. ocellicauda* are highlighted in yellow. C) Drops in the dS ratio along the gene correspond to the detected converted regions. D) A phylogenetic tree based on the longest converted region of the coding region i.e., exon5-6 (position 526 – 1074 of the genes) shows that the two *T. ocellicauda* sequences cluster together.

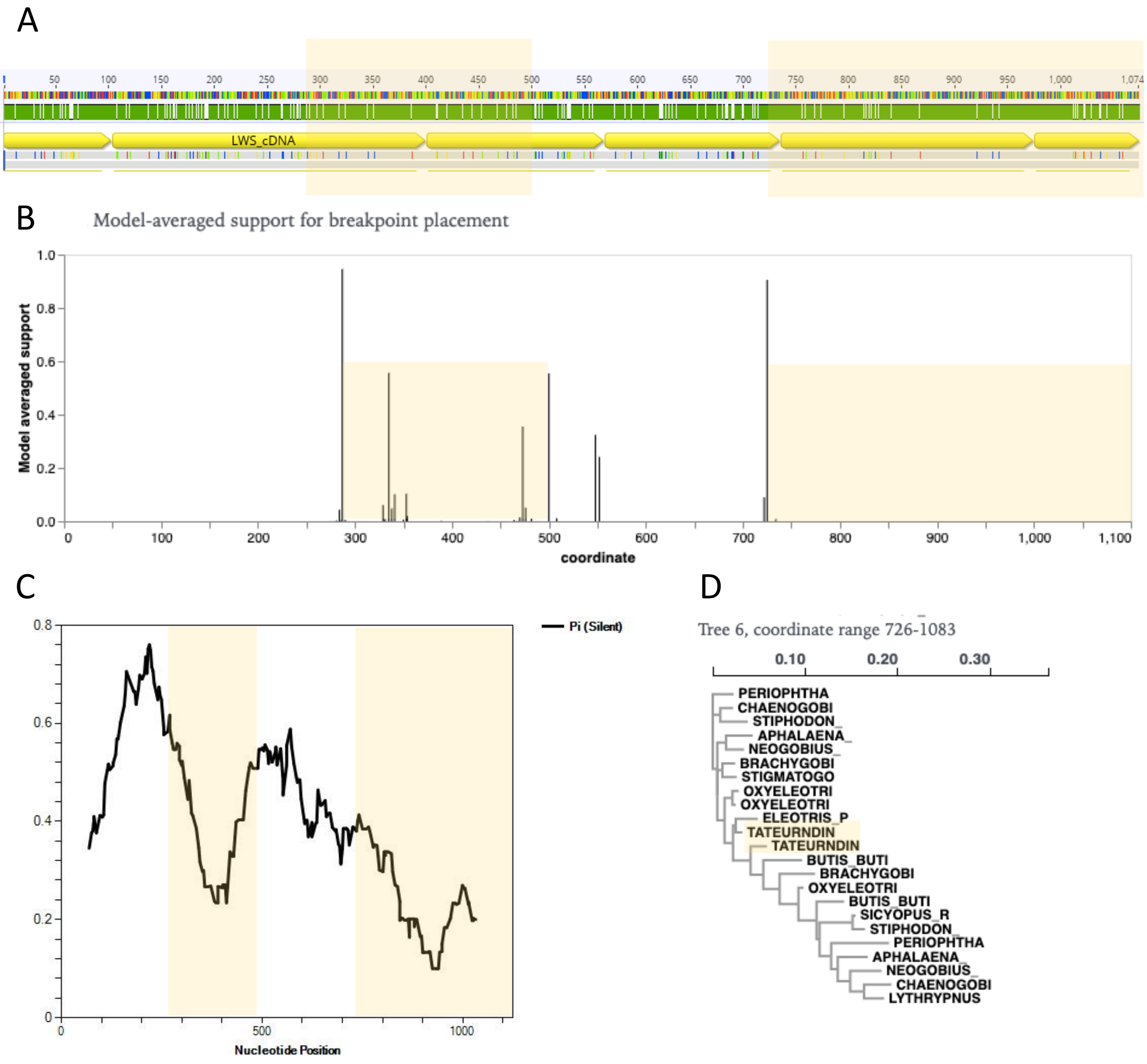

**Figure S7**: The miRNA detected in the upstream region of *LWS1* (all species), *LWS2* (*Megalops*, one copy) and *LWS3* (*Neogobius*).

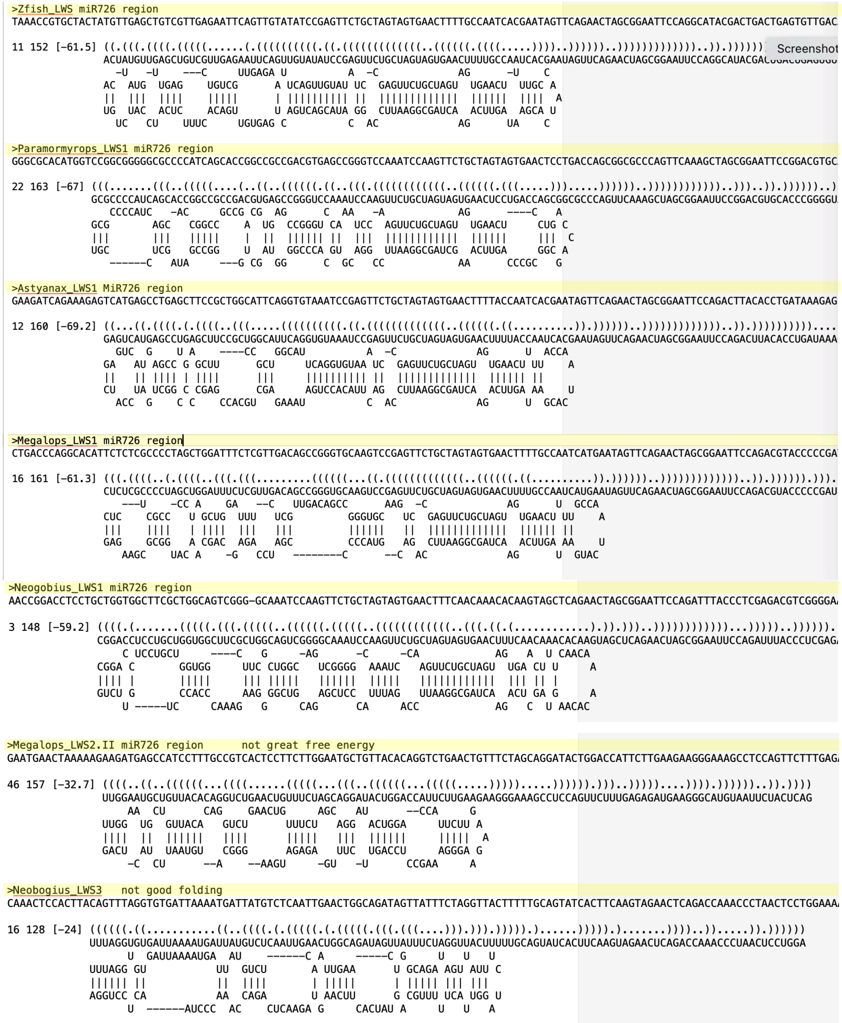


| | |
|---|---|
| **LWS1:** | |
| Zebrafish | -61.5 kcal/mole |
| *Paramormyrops* | -67 |
| *Astyanax* | -69.2 |
| *Megalops* | -61.3 |
| *Neogobius* | -59.2 |
| **LWS2**: | |
| Megalops | -32.7 |
| **LWS3**: | |
| Neogobius | -24 |

**Figure S8:** Structural models of LWS opsins in teleost fishes and in human

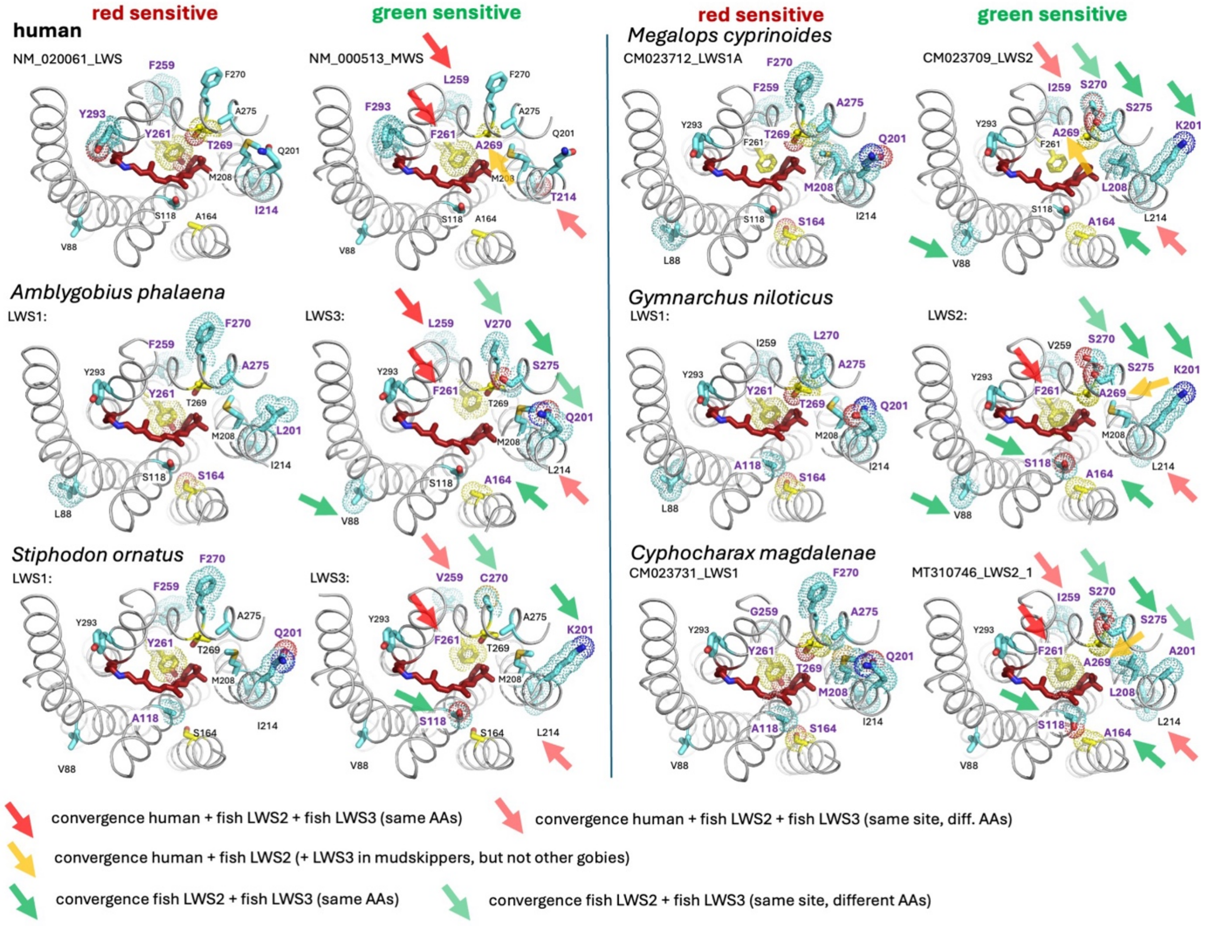

**Figure S9**: Double-labelling in-situ hybridisation of opsin mRNAs in retinal single cone (A-C) and double cone (D-F) photoreceptors of adult whitebarred goby, *Amblygobius phalaena*. **(A-C)** *SWS2B* (magenta) and *SWS2Aβ* (yellow) mRNAs were co-expressed in some single cones. **(D-F)** *LWS1* (magenta) and *LWS3* (yellow) mRNAs were expressed in opposite members of double cones. Scale bars: 10 µm.

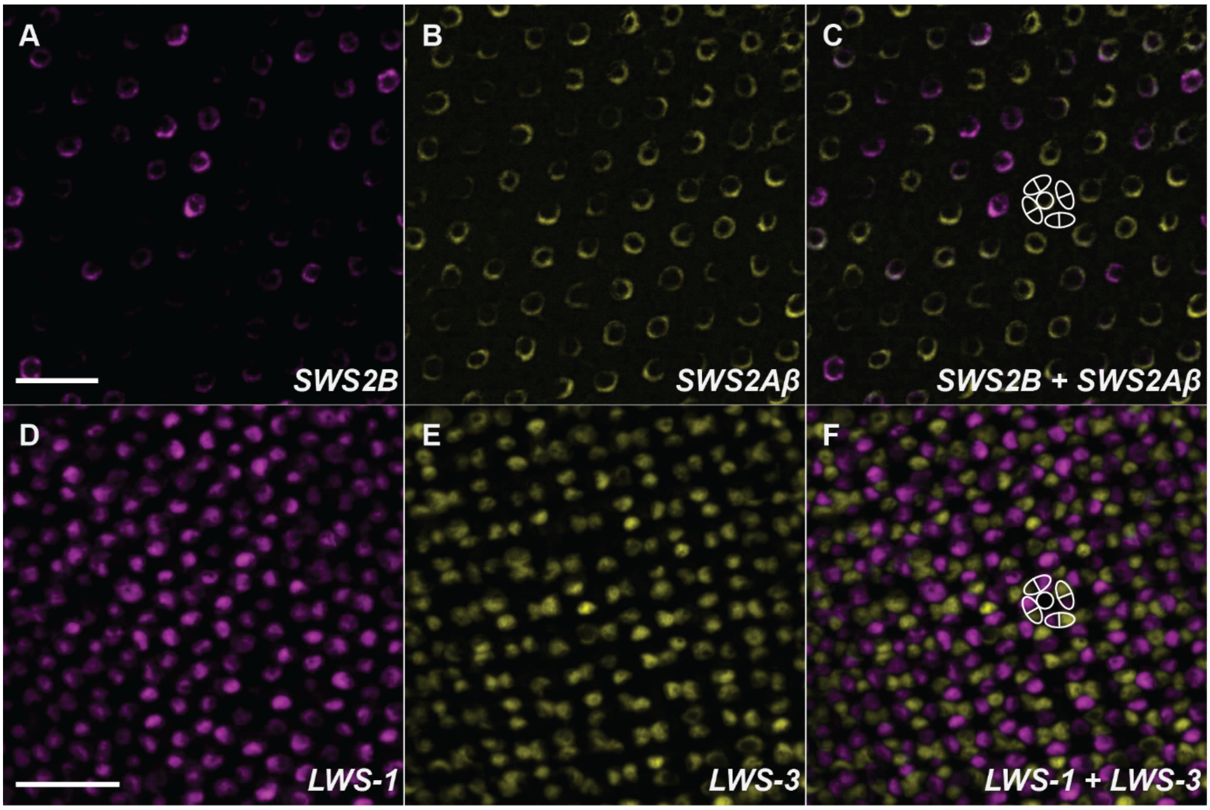
